# Single-cell RNA sequencing uncovers the excitatory/inhibitory synaptic unbalance in the retrosplenial cortex after peripheral nerve injury

**DOI:** 10.1101/2021.06.09.444962

**Authors:** Jing-Hua Wang, Cheng Wu, Yan-Na Lian, Zi-Yue Wang, Jia-jun Dong, Qin Wu, Li Liu, Li Sun, Wei Chen, Wenjuan Chen, Zhi Zhang, Min Zhuo, Xiang-Yao LI

## Abstract

Nerve injury in the somatosensory pathway may induce maladaptive changes at the transcriptional or protein level, contributing to the development and maintenance of neuropathic pain. In contrast to the retrosplenial cortex (RSC), which processes nociceptive information and exhibits structural and molecular changes after nerve injury, detailed transcriptional changes in the RSC are not yet known. Here we confirm the involvement of the RSC in regulating pain sensation and observe that the same peripheral stimulation activates more retrosplenial neurons after nerve injury; reducing the activities of *CaMKII*α^*+*^ splenial cells relieves peripheral pain hypersensitivity after nerve injury. Using a single-cell RNA sequencing (scRNA-seq) approach, we identified cell-type-specific gene expression changes after nerve injury, and the gene set enrichment analysis results revealed suppressed ion homeostasis in *CaMKII*α^+^ neurons. Furthermore, examination of the expression of genes encoding ligand-gated ion channels showed a decrease in *Gabar1a* but an increase in *Gria1* in *CaMKII*α^*+*^ neurons; consistently, we confirmed the unbalanced excitatory/inhibitory synaptic transmission by using the electrophysiological recording approach. Moreover, micro-infusion of 1-Naphthyl acetyl spermine in the RSC to reduce excitatory synaptic transmission alleviated peripheral pain hypersensitivity. Our data confirm the involvement of the RSC in pain regulation and provide information on cell type-dependent transcriptomic changes after nerve injury, which will contribute to the understanding of the mechanisms mediating neuropathic pain.

## INTRODUCTION

Neuropathic pain caused by injury to the somatosensory pathway affects approximately 8% of the world’s population; severely impairing the lives of patients ^[1]^. Unfortunately, the clinical management of neuropathic pain remains complex ^[2]^. Maladaptive plasticity occurs within the somatosensory pathways after nerve injury, from the dorsal root ganglion to the somatosensory cortex, which leads to pain hypersensitivity ^[3]^. However, the mechanisms that mediate these changes are complex. Multiple dysregulated molecular processes, such as gene transcription, protein translation and protein post-translational modifications, occur after nerve injury. Gene transcription after nerve injury contributes to irregular synaptic plasticity, abnormal neuronal excitability and altered interbrain region connectivity, which leads to irregular pain sensation ^[4–6]^. Knowing the transcriptional changes induced by nerve injury provides a novel window to understand the molecular mechanisms of neuropathic pain.

Peripheral nerve injury may induce extensive transcriptomic changes in different cell types. In the central nervous system (CNS), glial cells such as astrocytes, oligodendrocyte and microglia have unique roles in maintaining brain function. Moreover, numerous studies have shown that astrocytes and microglia undergo pathological changes that contribute to the development and maintenance of chronic pain ^[7,8]^. The recently developed single-cell RNA sequencing (scRNA-seq) approach is helpful in studying the heterogeneity of cell types. Using this approach, Xu Zhang’s group identified 10 types and 14 subordinate subtypes of somatosensory neurons in the dorsal root ganglion (DRG) of mice ^[9]^. The results revealed a distinct and sustained heterogeneity of transcriptomic responses to injury at a single neuronal level ^[10]^. Except for DRG neurons, the scRNA-seq approach was also applied to the spinal cord in the mouse neuropathic pain model to reveal novel regulators of pain hypersensitivity and provide new therapeutic targets for neuropathic pain ^[11]^. By screening differentially expressed genes in DRG of mice with spared nerve injury (SNI), activating transcription factor 3 (ATF3) was significantly increased after SNI stimulation, indicating that ATF3 plays an essential role in the progression of neuropathic pain and could be used as a candidate genetic marker for the prognostication and treatment of neuropathic pain ^[5]^. Jun, a proto-oncogene that acts as a transcription factor, has also been identified to interact strongly with other genes; it may serve as prognostic and predictive genes for neuropathic pain ^[12]^. Therefore, scRNA-seq provides a reliable and stable method to reveal cellular heterogeneity and candidate genetic markers in the progression of neuropathic pain caused by peripheral injuries.

The retrosplenial cortex (RSC) processes noxious information ^[13,14]^. RSC has been shown to reciprocally connect hippocampal formation and thalamus, regulate place navigation ^[15–17]^ and respond to nociceptive stimuli under physiological conditions ^[18]^. Furthermore, approximately 23% of RSC neurons responded to cutaneous nociceptive stimulations in anesthetized rabbits ^[13]^, and peripheral pain stimulation enhanced regional cerebral blood flow (rCBF) ^[19]^ and induced rigorous emotion-like behavior, followed by higher amounts of c-Fos-positive neurons in RSC of rats ^[20]^. In addition, spinal nerve ligation decreased metabolism in the RSC of mice ^[21]^. These observations indicate that metabolic and connectome changes occur in the RSC after nerve injury. Based on these studies, we propose that peripheral nerve injury may induce extensive transcriptional changes in the RSC; this may contribute to peripheral pain hypersensitivity. Given that both the anterior cingulate cortex and the RSC belong to the cingulate cortex ^[22]^, the molecular changes in the ACC have been broadly studied ^[23]^, whereas those in the RSC are unknown. Here, we investigated this point in the RSC of a mouse with nerve injury, by combining single-cell RNA sequencing approaches. We identified differentially expressed genes in different cell types. Our data provide a whole landscape of transcriptomic changes in the RSC induced by nerve injury, which will contribute to the understanding of the molecular and cellular mechanisms of neuropathic pain.

## Materials and methods

### Experimental animals

In this study, adult C57B L/6 mice and Gad67-GFP (from the Takeshi Kaneko laboratory of Kyoto University) (8 weeks, weight: 20–35 g) were used, and the animals were housed four or five per cage at constant room temperature (21 ± 1°C) and relative humidity (60 ± 5%) under a regular light/dark schedule (light from 7 am-7pm) with food and water being available ad libitum. Mice were acclimatized to laboratory conditions for approximately one week prior to behavioral tests and to the test environment for at least 15 min prior to the experiment. The Zhejiang University Animal Care and Use Committee approved all mouse protocols.

### Stereotaxic virus injection

AAV9-CaMKIIα-hM3Dq-mCherry, AAV9-CaMKIIα-hM4Di-mCherry, and AAV9-CaMKIIα-mCherry were obtained from Vigene Biosciences (Shandong, China). Stereotaxic injections of AAVs were performed and adapted ^[24]^. Briefly, mice were anesthetized with isoflurane (induction 4%, maintenance 1%) and the scalp was shaved, then cleaned with iodine (Triadine) and alcohol. The head of the mouse was fixed in a stereotaxic adapter mounted on a stereotaxic frame (Kopf model 962) and lubricant (artificial tears) was applied to the eyes. An incision was made over the skull and the surface was exposed. Two small holes were drilled above the RSC (AP: −2.30 mm, ML: ±0.25 mm, VD: −1.20 mm) and the dura was gently reflected. The virus was infused at a rate of 20 nl/min. After infusion, the needle was kept at the injection site for 10 min and then slowly withdrawn. The total volume of virus infused was based on its titer.

### Common peroneal nerve (CPN) ligation model

CPN ligation was performed as described previously ^[25]^. Briefly, mice were anesthetized with isoflurane (1-3%, as needed). The left CPN was slowly ligated between the anterior and posterior muscle groups with chromic gut suture 5-0 (Ethicon) until the appearance of finger twitching. The skin was sutured with 5-0 silk and cleaned with povidone iodine. The sham surgery was carried out in the same manner, but without ligating the nerves. All animals were kept in their home cages after surgery.

### Behavioral tests

#### Mechanical allodynia test

On an experimental day, the von Frey behavioral assay was performed according to the up-down algorithm described by Dixon ^[26]^. To determine evoked reflex responses to mechanical stimuli, animals were placed on a raised mesh grid and covered with a clear plastic box for containment. Calibrated von Frey filaments were applied to the middle of the plantar surface of each paw until the filaments were bent. Brisk withdrawal or paw flinching was considered as a positive response. Lifting of the paw due to normal locomotor behavior was ignored. In the absence of a response, the filament of the next greater force was applied. Following a response, the filament of the next lower force was applied. The tactile stimulus that produces a 50% likelihood of withdrawal was calculated and treated as the paw withdraw threshold (PWT).

#### Conditioned place preference/aversion

A conditioned place preference (CPP) test was performed, as described previously ^[27]^. Briefly, mice were preconditioned for three days: on the first two days, they were allowed to freely explore the chamber for 30 min; a video camera recorded the behaviors on the third day and we analyzed the time spent in each chamber; the next day, we injected the control solution being paired with a randomly chosen chamber in the morning, and the appropriate drug treatment being paired with the other chamber 4 h later (in the afternoon). 20 h after pairing in the afternoon, we placed mice in the CPP box with access to all chambers and analyzed their behavioral recordings for 15 min to understand chamber preference. The preference index was calculated as the time spent in the drug-paired chamber minus the time spent in the vehicle-paired chamber.

#### Novel object recognition

The novel object recognition test was adapted from a behavioral paradigm reported by Leger M. et al. ^[28]^. Briefly described, the mice were first placed in an open field chamber for 10 min to habituate. At the second day, two identical objects were placed in the diagonal of the chamber. One hour after CNO intraperitoneal injection, mice were allowed to explore two identical objects for 5 min. 24 h later, one of the objects was replaced with a new one of a different shape and the mice were allowed to freely explore for 5 min, and the time spent exploring the old and new objects was recorded, respectively. The percent of time spent in two objects was calculated to show the difference.

#### Elevated plus maze test

The elevated plus maze has been validated for assessing anxiolytic effects and defining brain regions and animals’ behavior associated with anxiety. After 30 min of CNO application, the mouse was placed at the junction of the closed and open arms, with its head facing the open arm. The video tracking system was then started and recorded for 5 min.

#### Open field test

White plastic boxes were used as open field chambers (dimensions: 45 × 45 × 45 cm^3^). Mice were placed individually in the center of the chamber and allowed to explore freely for 10 min. Locomotor and exploratory behaviors were recorded using ANY-maze software (Stoelting, Wood Dale, IL 60191, USA). The total distance traveled was used to evaluate locomotor activity.

#### Whole-cell patch-clamp recording

Coronal brain slices (300 μm) at the RSC level from sham-operated or CPN-ligated mice were prepared using standard methods. The slices were transferred to a submerged recovery chamber with oxygenated (95% O_2_ and 5% CO_2_) artificial cerebrospinal fluid (ACSF) containing (in mM) 124 NaCl, 2.5 KCl, 2 CaCl_2_, 1 MgSO_4_, 25 NaHCO_3_, 1 NaH_2_PO_4_, and 10 glucose at room temperature for at least 1 h. Experiments were performed in a recording chamber on the stage of a microscope equipped with infrared differential interference contrast optics for visualization. Spontaneous excitatory postsynaptic currents (sEPSCs) and Spontaneous inhibitory postsynaptic currents (sIPSCs) were recorded from layer II/III neurons of the RSC using an Axon 700B amplifier. sEPSCs were recorded in a voltage clamp model with membrane potentials at −70 mV. Recording micropipettes (3-5 MΩ) were filled with a solution containing (in mM) 124 K-gluconate, 5 NaCl, 1 MgCl_2_, 0.2 EGTA, 10 HEPES, 2 Mg-ATP, 0.1 Na_3_GTP, and 10 phosphocreatine disodium salt (adjusted to pH 7.2 with KOH). sIPSCs were recorded at 0 mV. Recording micropipettes (3-5 MΩ) were filled with a solution containing (in mM) 120 CsCl, 5 NaCl, 10 EGTA, 10 HEPES, 1MgCl_2_, 0.3 Na_2_-ATP, 3 Mg-ATP. (adjusted to pH 7.2 with KOH). The initial access resistance (15–30 MΩ) was monitored throughout the experiment. Data were discarded if the changes in access resistance were >15% during the experiment. Data were filtered at 1 kHz and digitized at 10 kHz.

#### Immunostaining

Mice were anesthetized with 1% sodium pentobarbital and perfused transcardially with 0.1◻M phosphate-buffered saline (PBS) followed by 4% paraformaldehyde (PFA) in PBS. Brains were post-fixed overnight at 4°C in PFA. Each brain was then dissected and further fixed in 4% PFA for an additional 24◻h, and then transferred to 15% sucrose in PB followed by a transfer to 30% sucrose until saturation. The brain was embedded in Tissue-Tek OCT compound, frozen in liquid nitrogen, and stored at −80°C before being cut into 25 μm coronal sections in a cryostat at –20°C (CM3050S, Leica). Free-floating sections were washed with PBS. For immunostaining, sections were incubated with blocking buffer (5% normal goat serum and 0.3% Triton X-100 in PBS) for 1 h at room temperature, and then incubated with primary antibody (c-Fos, 1:1000; #2250, Cell Signaling Technology, Danvers, USA) overnight at 4°C. Sections were washed in PBS and then incubated with the appropriate secondary antibody (goat anti-rabbit 488; 1937195, Life Technologies) for 2◻h at room temperature. Sections were re-washed in PBS for 3×10 min. After washing, they were mounted on coverslips using Fluoroshield Mounting Medium With DAPI (ab104139, Abcam, Cambridge, UK) for image collection.

### Cannulation and microinjection

Mice were anesthetized with ketamine (100 mg/kg body weight) and xylazine (8 mg/kg) by intraperitoneal (i.p.) injection and placed in a stereotactic frame. The scalp was shaved and cleaned with iodine and alcohol. An incision was made over the skull and the surface was exposed. Two small holes were drilled above the RSC and the dura was gently reflected. Guide cannulas were placed 2.25 mm behind bregma, 0.25 mm lateral to the midline, and 0.6 mm ventral to the surface of the skull.

For microinjection, mice were anesthetized with isoflurane inhalation in 100% oxygen at 0.5 L/min through a facemask. The dummy cannulas were removed and the microinjection cannula was inserted into the guide. A 30-gauge injection cannula was placed 0.5 mm below the end of the guide. The vehicle or Naspm (0.5 mL/side, 10 mM, MCE, Shanghai, China) was delivered bilaterally at a rate of 0.15 mL/min using a syringe driven by an infusion pump (ALC-IP600, Alcott, Shanghai, China). The volume delivered was confirmed by observing the movement of the meniscus down the length of the calibrated polyethylene (PE10) tubing. After delivery to each side of the brain, the injection cannula was left in place for 2 min to minimize back-flow along the guide. The cannula was then retracted and inserted into the opposite side of the brain. 30 min after microinjection, mechanical allodynia or thermal withdraw latency was tested.

#### Hargreaves test

The hargreaves test was adapted from a behavioral paradigm reported by Hargreaves K. et al. ^[29]^. Briefly described, the mouse was kept on the testing arena for 2 h and allowed to acclimate to the test environment 3 days prior to the test. The intensity of the light source was 16% and the cut-off time was 20 s. During the test, the radiant heat source was positioned underneath the animal and aimed at the plantar surface of the hind paw. The time required to withdraw from the heat stimulus was recorded as the withdrawal latency. The test was repeated so that each mouse was tested four times with the same paw.

#### Quantitative Real-time PCR Assay

Reverse transcription (RT) was performed using a PrimeScript® RT Reagent Kit with gDNA Eraser (RR047A, TaKaRa, Dalian, China) in a 20 μL reaction mixture containing 1 μg of total RNA from each individual sample. Primer pairs used to detect genes of interest are listed bellow. Quantitative real-time PCR was carried out in a total volume of 10 μL, with each tube containing 5 μL of SYBR Premix Ex Taq (Cat# 11201, Yeasen, Shanghai, China), 1 μL of RT product (40 ng) and 0.2 μL of primers (400 nM each). Three replicates were conducted for each sample. Reactions were run on a Roche LightCycler 480II at 95◻ for 3 min, followed by 40 cycles at 95◻ for 5 s and 60◻ for 15 s. We used the ratio between the gene of interest and β-actin to calculate the relative abundance of mRNA in each sample. Relative quantification of mRNA was calculated using the comparative CT (2^−ΔΔCT^) method ^[30]^.

The primer sequence for the examination of gene expressional changes using RT-qPCR

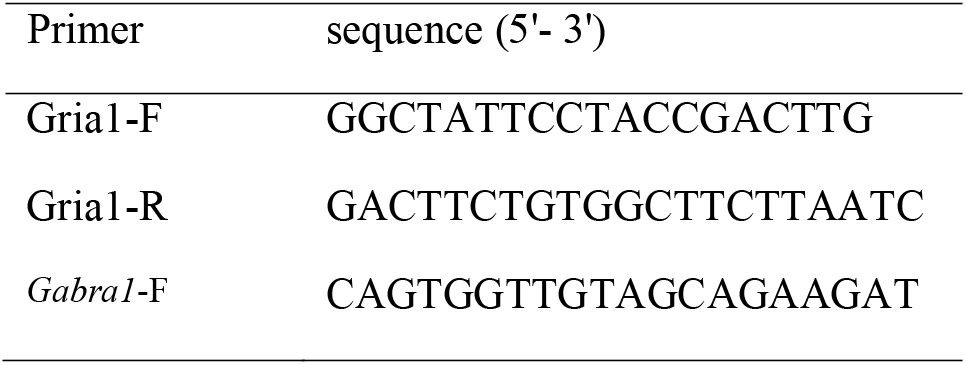
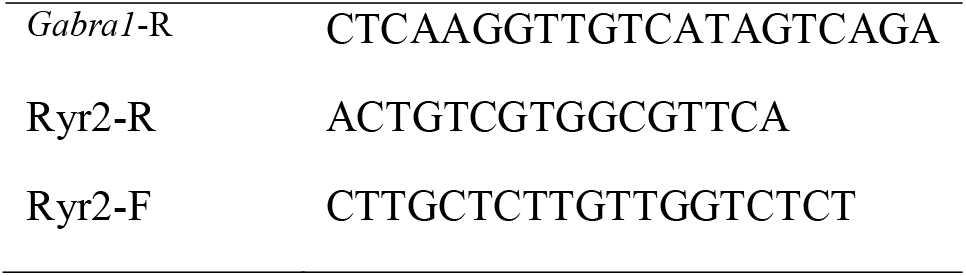

#### Cell isolation

Single cells were obtained according to a previous protocol ^[31]^. Individual adult male mice were anesthetized with isoflurane and decapitated. The brain was removed and the RSC was quickly dissected out. The RSCs from both mice were placed together and the tissue was cut into small pieces and transferred into a 1.5 ml microcentrifuge tube with 3 mg/ml isolation solution containing pronase (Sigma, Cat#P6911-1G), 1% BSA and 50 μg/ml DNaseI (Sigma, cat. no. D5025) in 1 ml Hibernate A (Invitrogen, cat. no. A1247501)/B27 (Invitrogen, cat. no. 17504) medium (HABG). The tissue and solution were mixed for 30 min at 37◻ in a horizontal shaker with 200 g. After incubation, the fabric was gently triturated with polished tips and single cells were released. Purified cells were obtained by a density gradient centrifugation at 800 g for 15 min and resuspended in 1xPBS (without calcium and magnesium) containing 1% BSA, followed by centrifugation at 200 g for 2 min. Single cells were concentrated and resuspended in the desired medium. Cell number and health status were measured by trypan blue staining.

#### RNA library preparation for high-throughput sequencing

Thousands of cells were partitioned into nanoliter-scale Gel Bead-In-Emulsions (GEMs) using 10x GemCode Technology, where cDNA generated from the same cell shares a common 10x Barcode. Upon dissolution of the single-cell 3′ gel bead in a GEM, primers containing an Illumina R1 sequence (read1 sequencing primer), a 16-bp 10x Barcode, a 10-bp randomer, and a poly-dT primer sequence were released and mixed with the cell lysate and Master Mix. After incubation of the GEMs, barcoded, full-length cDNA from poly-adenylated mRNA was generated. Then the GEMs were broken, and silane magnetic beads were used to remove excess biochemical reagents and primers. Prior to library construction, enzymatic fragmentation and size selection were used to optimize the cDNA amplicon size. P5, P7, a sample index and R2 (read 2 primer sequence) were added to each selected cDNA during end repair and adapter ligation. The P5 and P7 primers were used in Illumina bridge amplification of the cDNA (http://10xgenomics.com). Finally, the library was sequenced into 150-bp paired-end reads using Illumina HiSeq4000.

#### Data processing of scRNA-seq

Single-cell RNA-seq was analyzed similarly to a previous study ^[32]^. Described briefly, a cell ranger 2.0.1 (http://10xgenomics.com) was used to perform quality control and read counting of Ensemble genes with default parameters (v2.0.1) by mapping to the mm10 mouse genome. Poor quality cells were excluded after using the gene-cell data matrix generated by Cell Ranger software. We also used the DoubletFinder and deleted the double cells. We then used Seurat (v3.2.3) to integrate the scRNA-seq assay. Only cells expressing more than 200 genes and less than 7,000 genes were considered, and only genes expressed in at least three single cells (0.1% of the raw data) were included for further analysis. Cells expressing hemoglobin genes (*Hbm*, *Hba1*, *Hba2*, *Hbb*, *Hbd*, *Hbe1*, *Hbg1*, *Hbg2*, *Hbq1*, and *Hbz*) were excluded. Cells with more than 10% of mitochondrial genes were discarded. In total, 20,449 genes out of 43,275 single cells were used for subsequent analysis. The data were normalized to a total of l × 10^4^ molecules per cell for sequencing depth using the Seurat package. The canonical correlation analysis (CCA) from Seurat was used to correct the batches effect. The two groups of samples were first integrated using the FindIntegrationAnchors function and the IntegrateData function, and then the data were scaled using the ScaleDate function of Seurat (v3.2.3).

#### Identification of cell types and subtypes by dimensional reduction

On day 7 after CPN ligation or sham treatment, the present study conducted cell isolation and performed single-cell RNA sequencing experiments. Linear dimensional reduction was performed using the Seurat package (v3.2.3) ^[32]^. We selected 2000 highly variable genes as input for PCA. We then identified significant PCs based on the JackStrawPlot function. Strong PC1–PC50 was used for t-SNE to cluster cells using the FindClusters function with a resolution of 0.5. Further construction was carried out on the K-Nearest Neighbor (KNN) graph based on the Euclidean distance in PCA space and cells clustered by the Louvain algorithm. The clusters with less than 100 cells and only one group of cells were excluded. The study only included clusters with cells, thereby ultimately obtaining 31,146 single cells for further analysis (Ctrl: 7,965 and 11,296 cells; CPN: 5,966; and 5,919 cells). Clusters were identified by the expression of known cell-type markers. A combination of SingleR ^[33]^ results and expression of the following markers was used to annotate cell types: *Pdgfra*, *Cldn5, Cx3cr1, Gja1, Cldn11, Acta2 and Kcnj8* were used to classify oligodendrocyte precursor cells (OPCs), endothelial cell, microglia, astrocyte, oligodendrocyte, vascular smooth muscle cell (vSMC) and pericyte, respectively. The clusters that mainly expressed *CaMKII*α and *Thy1* were treated as neurons. In response to the inconsistency between the SingleR prediction result and the classic marker annotation, we identified cell types mainly by the classic marker and further confirmed by other markers.

#### Identification of differentially expressed genes

Characterized differentially expressed genes (DEGs) of each cluster were identified using the FindMarkers function (thresh.use = 0.25, test.use = “wilcox”) and the Seurat R package ^[34]^. A Wilcoxon rank-sum test (default) was used and genes with Log_2_FC > 0.58 and *P* < 0.05 were selected as featured marker genes.

For comparison between the sham and CPN groups, DEGs for each cell type were identified using the FindMarkers function (thresh.use = 0.10) and the Seurat R package. ^[34]^ To confirm the DEGs, we used three methods (wilcox, bimod and MAST) and compared the expressional difference, with Log_2_FC > 0.25 for genes, and adjusted *P* < 0.05 as DEGs. DEGs detected by at least two methods were treated as DEGs for that cell type. The UpSet plot of DEGs were plotted using the UpsetR package (v1.4.0) ^[35]^.

The markers of the different layers were identified by using the Allen Brain Atlas (ISH Data :: Allen Brain Atlas: Mouse Brain (brain-map.org)) ^[36]^. After careful inspection, we used *Gucy1a1* and *Ddit4l* as markers for layers II & III, and *Etv1* and *Slc6a7* as markers for layer V. We first isolated *CaMKII*α^+^ cells using the subset functions and then used the sample information to separate the cells from the sham-operated group or CPN group; in addition, we isolated *Gucy1a1^+^* or *Ddit4l^+^* cells as layer II & III cells and *Etv1*^+^ or *Slc6a7*^+^ as layer V cells, and then compared the different expression of genes using the FindMarkers function (thresh.use = 0.10) and the Seurat R package. ^[34]^ Similarly, we used three methods (wilcox, bimod and MAST) and compared the expressional difference, with Log_2_FC > 0.25 for genes, and adjusted *P* < 0.05 as DEGs. DEGs detected by at least two methods were treated as DEGs of that cell type.

Given that the DEGs from the three methods (wilcox, bimod and MAST) are quite similar, the DEGs from the bimod method were used to perform Gene Set Enrichment Analysis (GSEA) via the ClusterProfiler R package (V3.16.1) ^[37]^. DEGs were first collected using the FindMarkers function (min.pct = 0.10, logfc.threshold = 0.10, only.pos=FALSE, test.use = “bimod”), and the official symbols of the DEGs were changed to ENTREZID using the “bitr” function of Clusterprofiler R package, with the DEGs being sorted with ENTREZID by the averaged logFC and then the GSEA being executed using the “gseGO” function. We further filtered Go terms using the clusterProfiler::simplify function (cutoff = 0.6) and plotted the top20 GO terms using the plotGOgraph function. The first child terms for biological process (BP) or molecular function (MF) were also identified on the qucikGO website (https://www.ebi.ac.uk/QuickGO/).

We analyzed the Cell-Cell Communications of Con or CPN separately using CellChat R package ^[38]^ following the general protocol. Since the gene interaction database of CellChat is different from the iTALK database ^[39]^, we combined these two together. For the interaction inference, we used “computeCommunProb” to calculate the communication probability, and filterCommunication (min.cells = 100) to filter the communication probability. For the comparations between CPN and Con, extract the ligand-receptor pairs with upregulated ligands in CPN with ligand.logFC = 0.25, receptor.logFC = NULL, and extract the downregualted ligand-receptor pairs ligand.logFC = −0.25, receptor.logFC = NULL.

Correlations were calculated using the corr.test function from the psych package (2.1.6) based on the average expression of the anchor genes ^[40]^. All *p*-values were less than 0.01.

#### Cell sorting

Cell sorting was performed on a Sony SH800 cell sorter equipped with a 100 μm microfluidic sorting chip, and only YFP or mCherry positive cells were sorted directly into Eppendorf tubes containing 700 μl of cell lysis buffer for bulk sorts. Cells were identified based on the gating strategy. Performance of the cell sorter was checked before the experiments using Autosetup beads (Sony Biotechnology).

#### Data availability

The scRNA-seq data used in this study is deposited in the Genome Sequence Archive (CRA004603). The raw image files used in the figures that support the findings of this study are available from the corresponding authors upon reasonable request.

#### Data analysis

Off-line analysis of whole-cell patch-clamp data was performed using Clampfit 10. GraphPad Prism 8.0 was used to plot and fit the data. Statistical comparisons were made using Student’s *t-*test, one-way ANOVA or two-way ANOVA (Tukey’s test, Bonferroni test, or Sidak’s test was used for *post hoc* comparison) or Kruskal-Wallis test (Dunn’s Multiple Comparison Test was used for *post hoc* comparison). All data are expressed as mean ± SEM. In all cases, *P* < 0.05 was considered statistically significant.

## RESULTS

### Enhancing the activities of *CaMKII*α^*+*^ neurons in the RSC via chemogenetic approaches changes somatic sensation in naïve mice

Previous studies have shown that noxious information emerges in the retrosplenial cortex (RSC) ^[13]^, which indicates that the RSC is involved in pain regulation. Here, we evaluated this point in the regulation of pain sensation by manipulating the activities of the alpha subunit of calcium/calmodulin-dependent protein kinase type II (*CaMKII*α^*+*^) neurons. The human M3 muscarinic DREADD receptor coupled to Gq (hM3D) was expressed on *CaMKII*α^*+*^ neurons (CaMKIIα-hM3D-mCherry) in RSC via adeno-associated virus (AAV) and AAV9-CaMKIIα-mCherry as controls **(Figure 1A)**, and Clozapine-N-oxide (CNO, 10μM) binding to this receptor increased neuronal excitability. Three weeks after virus injection, the effects of CNO were examined using the whole-cell patch-clamp recording approach. As shown in **Figure 1B and C**, CNO significantly increased the number of action potentials (APs) in hM3D(Gq)-expressing neurons (**Figure 1C**). Behaviorally, the systemic application of CNO decreased the paw withdrawal thresholds (PWTs), and this effect lasted for less than 24 h (**Figure 1D and E**). Furthermore, the CNO shortened the thermal withdrawal latency (**Figure 1F**). Therefore, enhancing the activation of *CaMKII*α^*+*^ neurons in the RSC sensitized both mechanical and thermal sensation.

**Figure 1.**
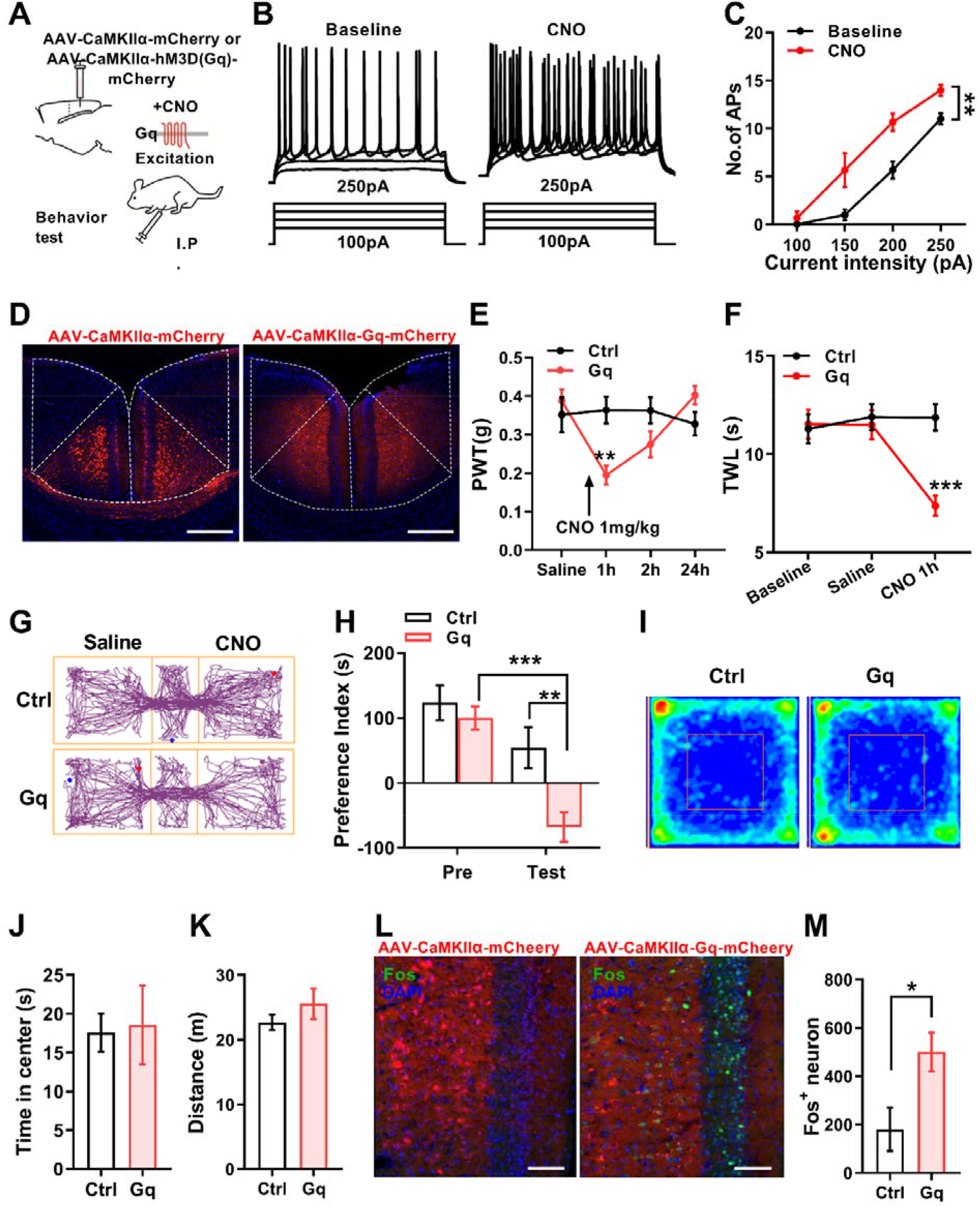
Activation of *CaMKII*α^*+*^ neurons in RSC decreases PWTs and induces conditioned place aversion. **(A)** Diagram showing the experimental design of hM3D expression. **(B)** Indication of traces showing that the application of CNO increased the number of APs in hM3D-expressing neurons. The raw traces show the individual voltage responses to 500 ms current pulses from 100 pA to 250 pA in 50 pA steps. **(C)** Summarized data showing increased excitability of hM3D-expressing neurons with CNO application (3 neurons from 2 mice, two-way ANOVA, Interaction, *F _(3, 16)_* = 2.57, *P* > 0.05; intensity, *F_(3, 16)_* = 75.18, *P* < 0.01; Baseline *vs.* CNO, *F_(1, 16)_* = 29.09, *P* < 0.01). **(D)** Representative images showing expression of control or Gq-mcherry in the RSC. **(E)** The application of CNO decreased the paw withdrawal thresholds (PWT) in Gq-expressing mice and recovered after 24 h (Two-way ANOVA, Interaction, *F _(3, 92)_* = 6.39, *P* < 0.01; time, *F_(3, 92)_* = 3.69, *P* < 0.05; Ctrl *vs.* Gq, *F_(1, 23)_* = 2.19, *P* > 0.05; Sidak’s multiple comparisons test, Ctrl *vs.* Gq (1 h): *P* < 0.01; Ctrl *vs.* Gq (2 h): *P* > 0.05, n = 10 for Ctrl, n =15 for Gq). **(F)** The application of CNO extended the paw withdrawal latency (PWL) in Gq-expressing mice (Two-way ANOVA, Interaction, *F _(2, 54)_* = 8.26, *P* < 0.01; time, *F _(2, 54)_* = 6.33, *P* < 0.01; Ctrl *vs.* Gq, *F_(1, 27)_* = 5.63, *P* <0.05; Sidak’s multiple comparisons test, Ctrl *vs.* Gq (1 h): *P* < 0.01; n = 16 for Ctrl, n =13 for Gq). **(G)** Representative traveling traces of the Ctrl and Gq group of mice in the conditioned place aversion test. **(H)** Place aversion induced by CNO application in the Gq group (Two-way ANOVA, Interaction, *F_(1,19)_* = 5.28, *P* < 0.05; time, *F_(1,19)_* = 30.41, *P* < 0.01; Ctrl vs. Gq, *F_(1,19)_* = 6.82, *P* < 0.05, Ctrl, n = 10, Gq, n = 11. Sidak’s multiple comparisons test, Ctrl *vs.* Gq (Test): *P* < 0.01; Pre *vs.* Test (Gq): *P* < 0.001). **(I)** Heat map diagram showing the traveling traces of both groups in the open field test. **(J)** Application of CNO did not change the timing of the central area in the open field test (Unpaired *t*-test, Unpaired *t*-test, n = 10 for Ctrl, n =6 for Gq, *P* > 0.05). **(K)** In the open field test, mice in the Gq group traveled a similar distance compared to Ctrl after CNO application (Unpaired *t*-test, n = 10 for Ctrl, n =6 for Gq, *P* >0.05). **(L)** Representative images of c-Fos expression induced by CNO injection in the RSC of naïve mice (Scale bar, 200 μm). **(M)** Application of CNO increased c-Fos expression in the Gq group (n = 5, Unpaired *t*-test, *P* <0.05).

Given that the neuronal activities of RSC are necessary for the aversive memory ^[41]^, does activation of *CaMKII*α^*+*^ neurons also elicit aversion? This point was examined by using a conditioned place behavioral approach (**Figure 1G**). In the pre-conditioning phase, mice with or without Gq expression spent equal amounts of time exploring the chambers **(Figure 1H**). While in the testing session, mice with Gq spent less time in the CNO paired chamber unlike mice with the control virus (**Figure 1H**). These data suggest that activation of *CaMKII*α^*+*^ neurons in RSC is sufficient to induce aversion. Furthermore, the CNO application did not alter the time spent in the central zone (**Figure 1I and J**) and traveling distance (**Figure 1K**) in the open field test, indicating that activating RSC *CaMKII*α^*+*^ neurons does not affect anxiety. In addition, we examined the expression of c-Fos (an immediate-early gene) and observed that CNO application increased the number of c-Fos positive cells in the RSC of mice with Gq expression (**Figure 1L and M**). In summary, activation of *CaMKII*α^*+*^ neurons in RSC via the chemogenetic approach sensitizes the mechanical sensation and induces aversion in naïve mice.

### CPN ligation altered neuronal activities in RSC contributing to the pain hypersensitivity

Peripheral nerve injury caused structural and metabolic changes in the RSC ^[42]^, indicating abnormal neuronal activities. To confirm this assumption, using a mouse model of neuropathic pain with ligation of the common peroneal nerve (CPN) ^[41]^, the expression of c-Fos in the RSC after mechanical stimulation with von Frey filaments (0.4 g) was examined. As shown in **Figure 2A** and **B**, a higher amount of c-Fos positive neurons was observed after mechanical stimulation on day 7 after CPN ligation. The anterior cingulate cortex (ACC) is vital for pain regulation ^[23]^; we also examined c-Fos expression in the ACC as a positive control and observed similar changes, but not in the hippocampus (**Figure 2A and D**). Besides, increased c-Fos was detected in almost all RSC layers (**Figure 2E**). Furthermore, about 92% of c-Fos positive neurons expressed *CaMKII*α (**Figure 2B and F**). In contrast, nearly no GABAergic neurons expressed c-Fos after mechanical stimulation (**Figure 2C**). These data suggest that mechanical stimulation with the same intensity activated more *CaMKII*α^*+*^ neurons in the RSC of CPN-ligated mice compared to the control group.

**Figure 2.**
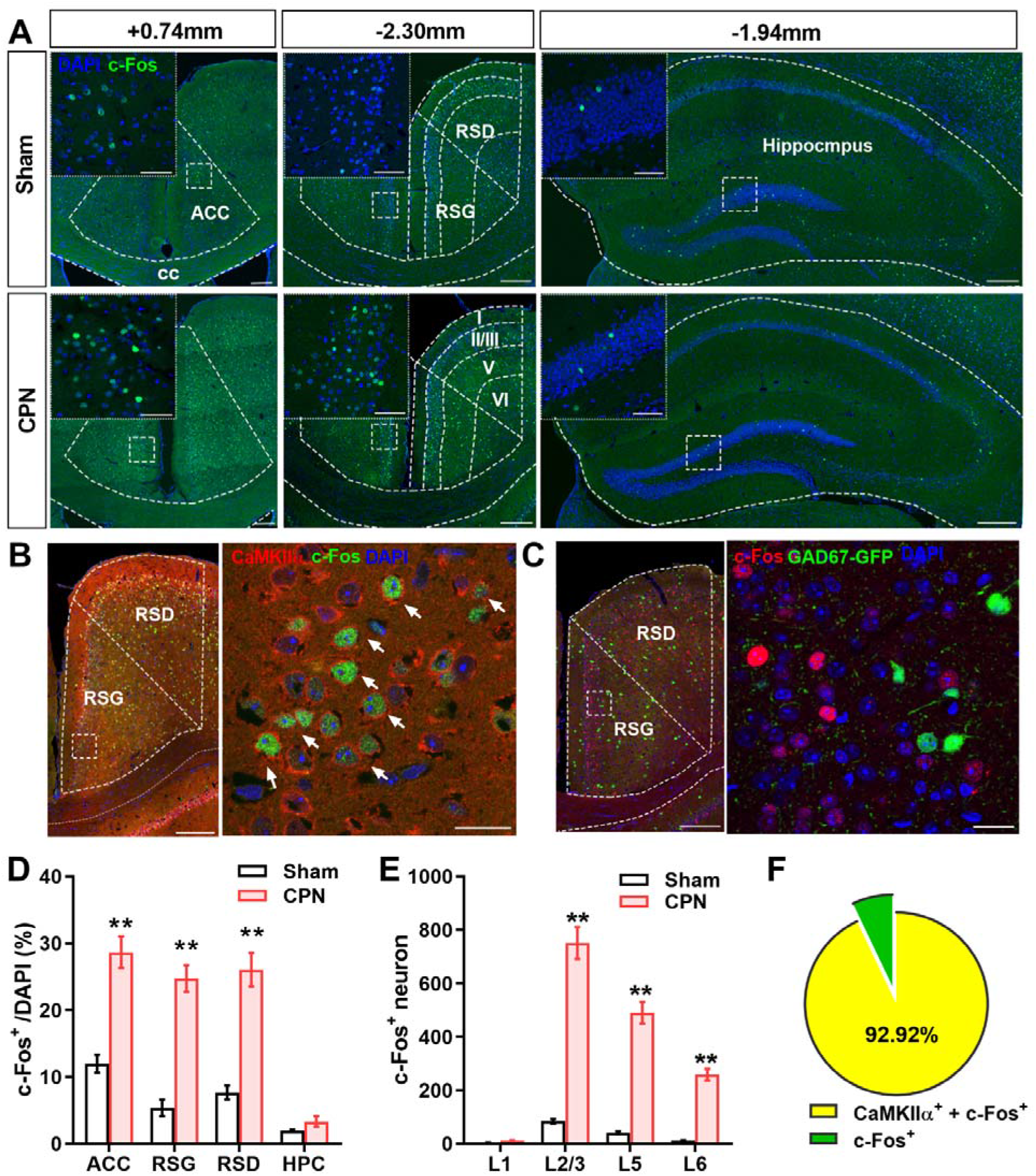
Mechanical stimulation activating more neurons in the RSC of CPN-ligated mice. **(A)** Examples showing the expression of c-Fos in different brain regions of the Sham or CPN group. Inset, region magnification (Bottom Scale bar, 200 μm; top left: 50 μm). **(B)** Example showing the c-Fos and CaMKIIα co-immunostaining in RSC (Left panel, scale bar, 200 μm; right panel, scale bar, 20 μm). **(C)** Example of imaging showing very few GABAergic neurons (GAD67) expressing c-Fos after mechanical stimulation. **(D)** Quantification of c-Fos positive cells within the ACC, RSC, and hippocampus (Hip), with more c-Fos positive cells in the ACC and RSC, but not in the hippocampus of CPN-ligated mice (Two-way ANOVA, interaction, *F_(3,31)_* = 32.19, *P* <0.01; regions, *F_(3,31)_* = 79.95, *P* <0.01; sham *vs.* CPN, *F_(1,31)_* = 311.90, *P* <0.01, Sham: n = 5, CPN: n = 5, ** indicates *P* < 0.01). **(E)** Summarized data showing more c-Fos^+^ cells in layer II-VI of RSC (Two-way ANOVA, n = 11 for Sham, n = 11 for CPN, Sidak’s multiple comparisons test, Sham *vs.* CPN (L2/3, L5,L6): *P* < 0.01.) **(F)** About 93.0% of c-Fos^+^ cells were CaMKIIα*^+^* cells.

Do neuronal activities in the RSC contribute to pain hypersensitivity after nerve injury? We selectively inactivated CaMKIIα-positive neurons and examined mechanical allodynia effects in mice. The human M4 muscarinic DREADD receptor coupled to Gi (hM4D) was expressed in the RSC (**Figure 3A and D**). Unlike hM3D, CNO bound to hM4D inhibits neuronal activities. We delivered the AAV9-CaMKIIα-hM4D-mCherry to the RSC. CNO application did decrease the number of firing APs examined via the whole-cell patch-clamp recording approach (**Figure 3B and C**). In animals with control or hM4D expression, we performed CPN ligation, which reduced the PWTs of both groups (**Figure 3E**). CNO application increased the PWTs of mice with Gi expression, but not of the control virus (**Figure 3E**). Similarly, CNO also extended thermal withdrawal latency in Gi-expressing mice with CPN ligation (**Figure 3F**). However, application of CNO to mice with CPN ligation did not alter the place preference for the CNO-paired chamber (**Figure 3G and H**), indicating that inhibition of the RSC did not alter the emotions of nerve injured mice. In summary, inhibiting the neuronal activities in the RSC changes the peripheral mechanical and thermal sensitivities after nerve injury.

**Figure 3.**
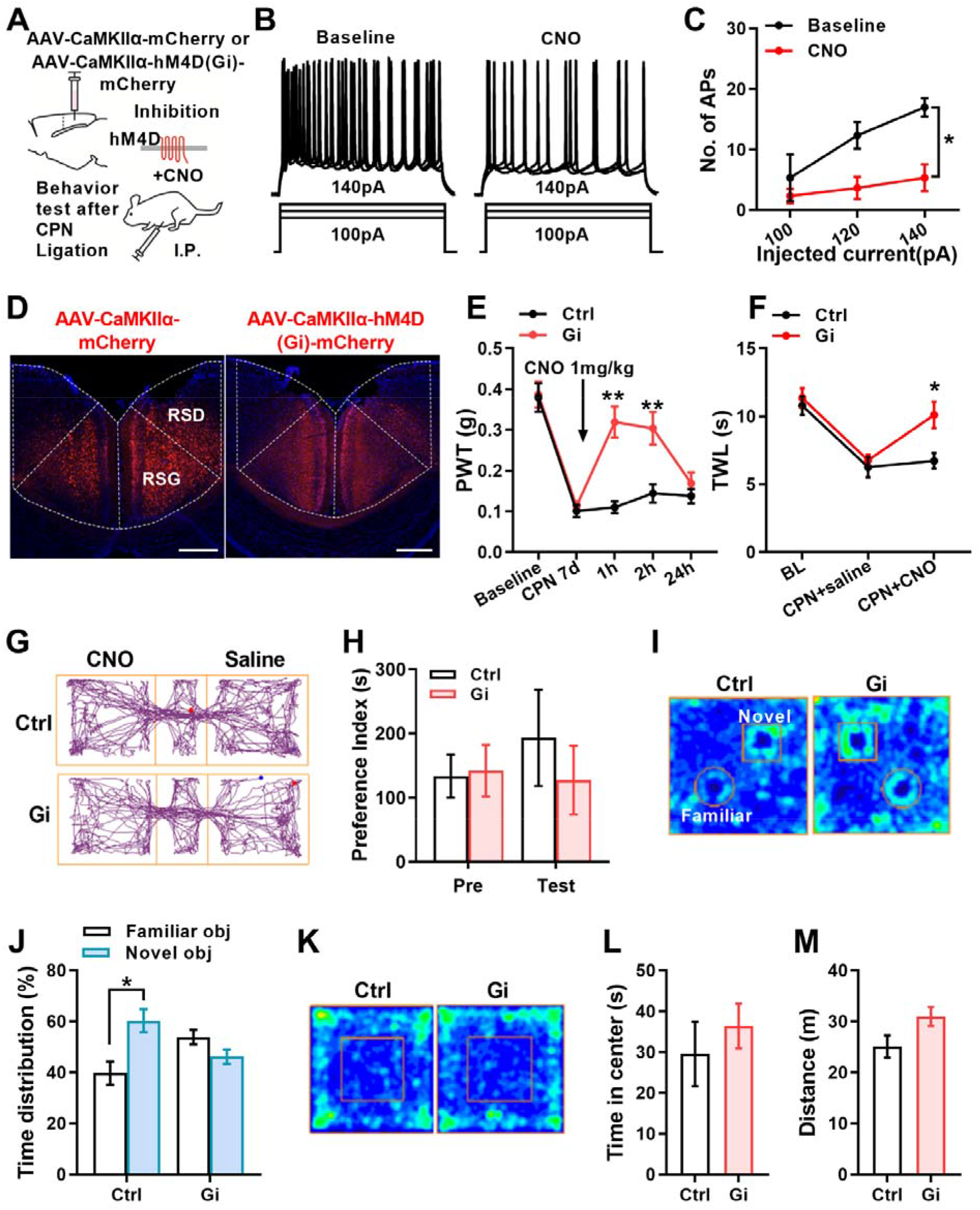
Inactivation of *CaMKII*α^*+*^ neurons in the RSC increases PWTs and extended TWL. **(A)** Diagram showing the experimental design of hM4D(Gi) expression. **(B)** Indication of traces showing that the application of CNO decreased the number of APs in Gi-expressing neurons. The raw traces show the individual voltage responses to 500 ms current pulses from 100 pA to 140 pA in steps of 20 pA. **(C)** Summarized data showing increased excitability of Gi–expressing neurons with CNO application (two-way ANOVA, interaction, *F_(2,8)_* = 2.92, *P* > 0.05; currents, *F_(1.1,4.4)_* = 8.16, *P* < 0.05; baseline *vs.* CNO, *F_(1,4)_* = 9.92, *P* < 0.05, 3 neurons from 2 mice) **(D)** Representative images showing expression of control or hM4D-mcherry in the RSC (Scale bar, 200 μm). **(E)** CPN ligation decreased PWTs in the Ctrl and Gi groups, and CNO application (i.p.) significantly increased PWTs in the Gi group with complete recovery after 24 h, having no effect on the Ctrl group (Two-way ANOVA, interaction, *F_(4,116)_* = 6.72, *P* < 0.01; time, *F_(4,116)_* = 33.51, *P* < 0.001; Ctrl *vs.* Gi, *F_(1,29)_* = 15.29, *P* < 0.01, n = 15 for Ctrl and n = 16 for Gi. Sidak’s multiple comparisons test, Ctrl *vs.* Gi (1h, 2h): *P* < 0.01). **(F)** CPN ligation shorted PWL in both groups, and CNO application (i.p.) significantly increased PWL in the Gi group (Two-way ANOVA, interaction, *F_(2,30)_* = 3.15, *P* > 0.05; time, *F_(1.96,29.37)_* = 24.44, *P* < 0.01; Ctrl *vs.* Gi, *F_(1,15)_* = 5.50, *P* < 0.05, n = 9 for Ctrl and n = 8 for Gi. Sidak’s multiple comparisons test, Ctrl *vs.* Gi (1 h): *P* < 0.05). **(G)** Representative traveling traces of the Ctrl and Gi group mice in the conditioned place preference test. **(H)** Application of CNO in the CPP test does not induce place preference in Gi group of mice (Two-way ANOVA, Interaction, *F_(1,9)_* = 0.91, *P* > 0.05; time, *F_(1,9)_* = 0.33, *P* > 0.05; Ctrl *vs.* Gi, *F_(1,9)_* = 0.21, *P* > 0.05, Ctrl, n = 5, Gi, n = 6). **(I)** Heat map diagram showing the traveling traces in the novel object recognition test. **(J)** Mice in the Gi group spent similar amounts of time exploring familiar and novel objects, whereas mice in the Ctrl group spent longer exploring novel objects (Two-way ANOVA, Interaction, *F_(1,20)_* = 13.67, *P* < 0.01; Ctrl *vs.* Gi, *F_(1,20)_* = 0, *P* > 0.05; Familiar *vs.* Novel, *F_(1,20)_* = 2.89, *P* > 0.05, n = 6 for both groups; Sidak’s multiple comparisons test, Familiar *vs.* Novel (Ctrl): *P* < 0.01). **(K)** Heat map diagram showing the traveling traces of both groups in the open field test. **(L)** In the open field test, mice in the Gi group moved a similar distance compared to Ctrl 1 h after the application of CNO (Unpaired *t*-test, n = 4 for Ctrl, n = 6 for Gi, *P* > 0.05). **(M)** In the open field test, mice in the Gi and Ctrl groups spent similar time in the central zone 1 h after the application of CNO (Unpaired *t*-test, n = 4 for Ctrl, n = 6 for Gi, *P* > 0.05).

We also expressed Gi in the RSC and performed the novel object recognition test (NOR). We found that applying CNO to mice carrying the control virus did not affect its performance in NOR and that the mice took longer to study the new object during the test period. However, mice with Gi expression spent the same amount of time on both familiar and novel objects (**Figure 3I and J**), indicating the necessity of the RSC for object memory. Although inhibition of CaMKIIα positive neurons in the RSC did not alter performance in the open field (**Figure 3 K-M**). Therefore, inhibition of neuronal activities in the RSC affects object memory.

### The single-cell landscape of RSC cells

In examining the cell type-dependent transcriptomic changes induced by CPN ligation in RSC, we first isolated RSC cells and analyzed the transcriptomic characteristics of retrosplenial cells using a single-cell RNA-sequencing (scRNA-seq) approach (**Figure 4A**). Then, we compared the differences in gene expression between mice with or without CPN. Since we intended to analyze the differences between the two groups, we included only clusters containing cells and eventually obtained 31,146 single cells for further analysis (Ctrl: 7,965 and 11,296 cells; CPN: 5,966; and 5,919 cells) belonging to seven types of cells (**Figure 4B, Figure S1A**) based on the expression of classical cell type markers (**Figure S1B**), including endothelial cells (ECs), microglia (Micro), oligodendrocyte precursor cells (OPC), oligodendrocytes (Oligo), astrocytes (Astro), *CaMKII*α^*+*^ neurons with high expression of *CaMKII*α, and vascular smooth muscle cells (vSMC). In our assay, we did not include GABAergic neurons. Then, using Seurat’s FindAllMarkers function, we identified the featured differentially expression genes (fDEGs) for each cell type (**Figure S1C**, Table S1).

**Figure 4.**
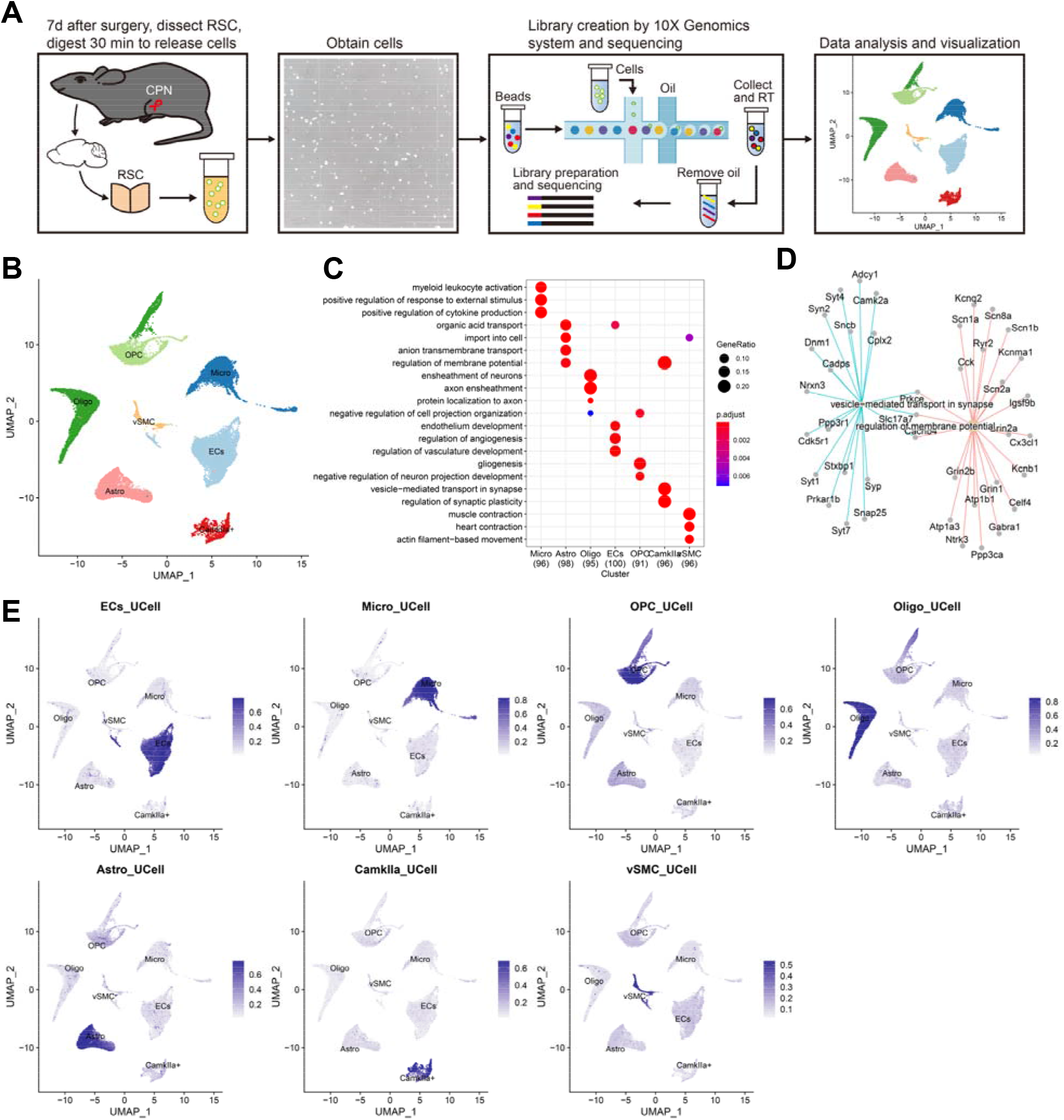
Transcriptomic profiles of single cells in the RSC. (A) Overview of the experimental workflow for single-cell RNA sequencing. (B) *Umap* embedding projection of seven cell types from the RSC of adult mice. (C) Dot plot showing the top3 GO terms of the biological process on the top 100 fDEGs. (D) Networks showing the BP terms “vesicle-mediated transport in synapse” and “regulation of membrane potential” and the enriched fDEGs of *CaMKII*α^+^ cells. (E) UCell score distribution in UMAP space for fDEGs signatures that are enriched in top3 BP terms evaluated using UCell (the fDEGs list is in the **Table S2**).

Next, a gene ontology (GO) analysis was performed on the top100 fDEGs, and different enriched biological process (BP) terms (**Figure 4C** and **Table S2, Figure S1D**) and cell type molecular function (MF) terms (**Figure S2A, Table S3**) were examined. For example, the top 3 BP terms on the Micro include myeloid leukocyte activation, positive regulation in response to external stimuli, and positive regulation of cytokine production, covering 29 fDEGs. For the Top3 MF terms, Micro reaches G protein-coupled purinergic nucleotide receptor activity, which includes genes *P2ry12*, *Gpr34*, and *P2ry13*, proteoglycan binding, and ribosomal related functions (**Figure S2B & C**). Consistent with the top 3 BP terms of Astro, which are organic acid transports imported into cell and associated with energy supporting functions, the organic acid transport terms were also detected in ECs, indicating cooperation between the two cell types. In MF analysis, the fDEGs of Astro reached transmembrane transport of sodium, carboxylic acids and organic anions. The transmembrane anion transport process may regulate ion homeostasis, similar to how some fDEGs of Astro regulate membrane potential in *CaMKII*α^+^ cells; therefore, regulation of membrane potential was detected in both Astro and *CaMKII*α^+^ cells. The BP terms for Oligo are axon ensheathment and protein-to-axon localization, both of which are associated with myelin sheath functions, and consistently, MF on Oligo is enriched to structural constituents of myelin sheaths due to the expression of Proteolipid Protein 1 (Plp1), myelin and lymphocyte protein (Mal), myelin-associated oligodendrocyte basic protein (Mobp), myelin basic protein (Mbp), and plasmolipin (Pllp). ECs and vSMC are part of the blood vessels, with fDEGs of ECs enriched in endothelial development and vasculature development, and fDEGs of vSMC related to muscle contraction and cardiac contraction, which are processes that may be mediated primarily by a consistent actin filament-based movement. The MF of fDEGs from ECs was enriched in cell adhesion-related binding and extracellular matrix, and these may be related to the formation of the blood–brain barrier. As a major cellular component of information processing, the fDEGs of *CaMKII*α^+^ cells were enriched in the regulation of membrane potentials, synaptic transmission and synaptic plasticity (**Figure 4C**); moreover, MF analysis indicated that its fDEGs were enriched to metal ion transmembrane transport, calmodulin binding, and glutamate receptor binding (**Figure S2B**). The example in **Figure 4D** shows the connection between fDEGs and BP terms: vesicle-mediated transport in the synapse and regulation of membrane potential. Other fDEGs and the related top 3 BP terms or top3 MF terms are shown in **Figure 1D** or **Figure S2C**, respectively. We further examined the dominant expression of these fDEGs by using the Ucell R package, and in line with the enriched BP terms, the Ucell score of fDEGs with top3 BP terms (**Figure 4E**) or MF terms (**Figure S2D**) indicates a cell-types dependent expression pattern.

### CPN ligation induces transcriptomic changes in the RSC

We next investigated the differential expression of the gene in the two groups. Cells from the Con (without CPN ligation) and CPN groups were projected into the uniform manifold approximation and projection (UMAP) for dimension reduction based on transcriptomic characteristics (**Figure 5A**). The differences in expressed genes on these cells were analyzed, and ECs, Micro, OPC and Oligo had higher correlation scores between the Con and CPN groups based on the average expression of the genes. In contrast, *CaMKII*α^+^ neurons and vSMC had lower correlation scores (**Figure 5B**), suggesting that CPN ligation may induce more dramatic changes in these cells. A comparison of gene expression differences between the Con and CPN groups was next performed via the FindMarkers function of Seurat (**Table S4**). **Figure 5C** presents the differentially expressed genes (DEGs) detected on *CaMKII*α^+^ neurons. Interestingly, some DEGs in *CaMKII*α^*+*^ neurons also exhibit altered expression in other cell types (**Figure 5D**), for example: this suggests that the expression of some DEGs is impaired in multiple cell types. Further analysis showed that about 22% (Common/Unique:77/281, **Table S4**) upregulated DEGs and 35% (Common/Unique: 115/211, **Table S4**) downregulated DEGs were detected in at least two cell types (**Figure 5E and F**), indicating that CPN ligation also induces cell type-dependent transcriptomic responses.

**Figure 5.**
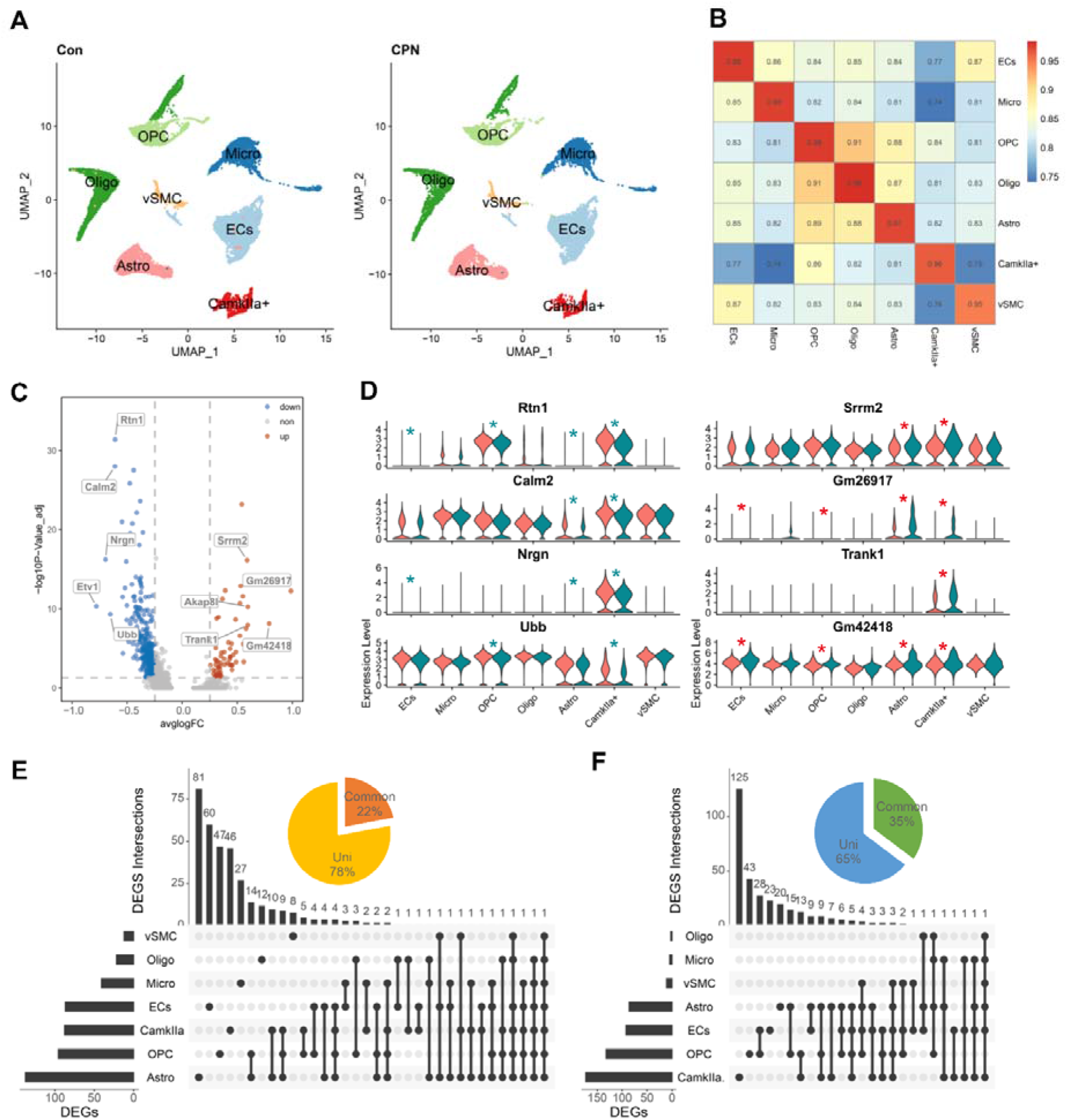
CPN ligation changes the transcriptional profile of cells in the RSC. (A) *t*-SNE embedding projections of cells from RSC of mice without (Con) or with CPN ligation (CPN). (B) Heatmap showing the correlation between the two cell types. (C) Volcano plots showing the DEGs between the Con and CPN groups detected in *CaMKII*α^*+*^ neurons. (D) Violin plot showing the expression of the DEGs with larger absolute long_10_P-Values in *CaMKII*α^*+*^ neurons. Sham group was painted in salmon, and CPN group was painted in dark cyan. “*” in red indicates upregulation while “*” in blue indicates downregulation. (E) Upset plot showing the distributions of upregulated DEGs among cell types. Inserted pie graph displayed the percent of common or cell-type specific DEGs. (F) Upset plot showing the distributions of downregulated DEGs among cell types. Inserted pie graph displayed the percent of common or cell-type specific DEGs.

### The cell type-dependent functional impairments

Gene set enrichment analysis (GSEA) indicates either activation or suppression effects via positive or negative normalized enrichment scores (NES), respectively (**Figure 6A**). More importantly, it also identifies the core enriched genes for the relevant terms. We performed a GSEA on the DEGs of each cell type and identified the different levels of GO terms. **Figure 6B** shows the ancestor charts for the GO term (GO:0007155) cell adhesion; this term is the final child terms for the cellular process (GO:0009987) and biological adhesion (GO:0022610), both of which are the first level of child terms for the biological process (GO:0008150). For significant final child terms (e.g., cell adhesion), we will statistically count the first level child terms of BP (e.g., biological adhesion). The normalized enrichment scores (NES) for each final child terms are shown in **Figure 6C and Table S5**. As shown in **Figure 6C**, forty-nine final child BP terms were detected, which belong to seven first level BP terms: including biological adhesion (BA, 2 terms), biological regulation (BR, 6 terms), cellular processes (CR, 7 terms), developmental processes (DP, 5 terms), localization (Lo, 5 terms), metabolic processes (MP, 16 terms), and response to stimuli (RtS, 8 terms). In the same way, we examined the impaired final child terms of molecular function (**Figure 6D, Table S6**) for each cell type and detected 46 final child MF terms belonging to 8 first-level MF terms, including antioxidant activity (AA), binding, catalytic activity, molecular function regulator (MFR, 2 terms), molecular transducer activity (MTA, 1 term), structural molecule activity (MA), transcriptional regulatory activity (TRA, 2 terms) and transporter activity (TA, 2 terms).

**Figure 6.**
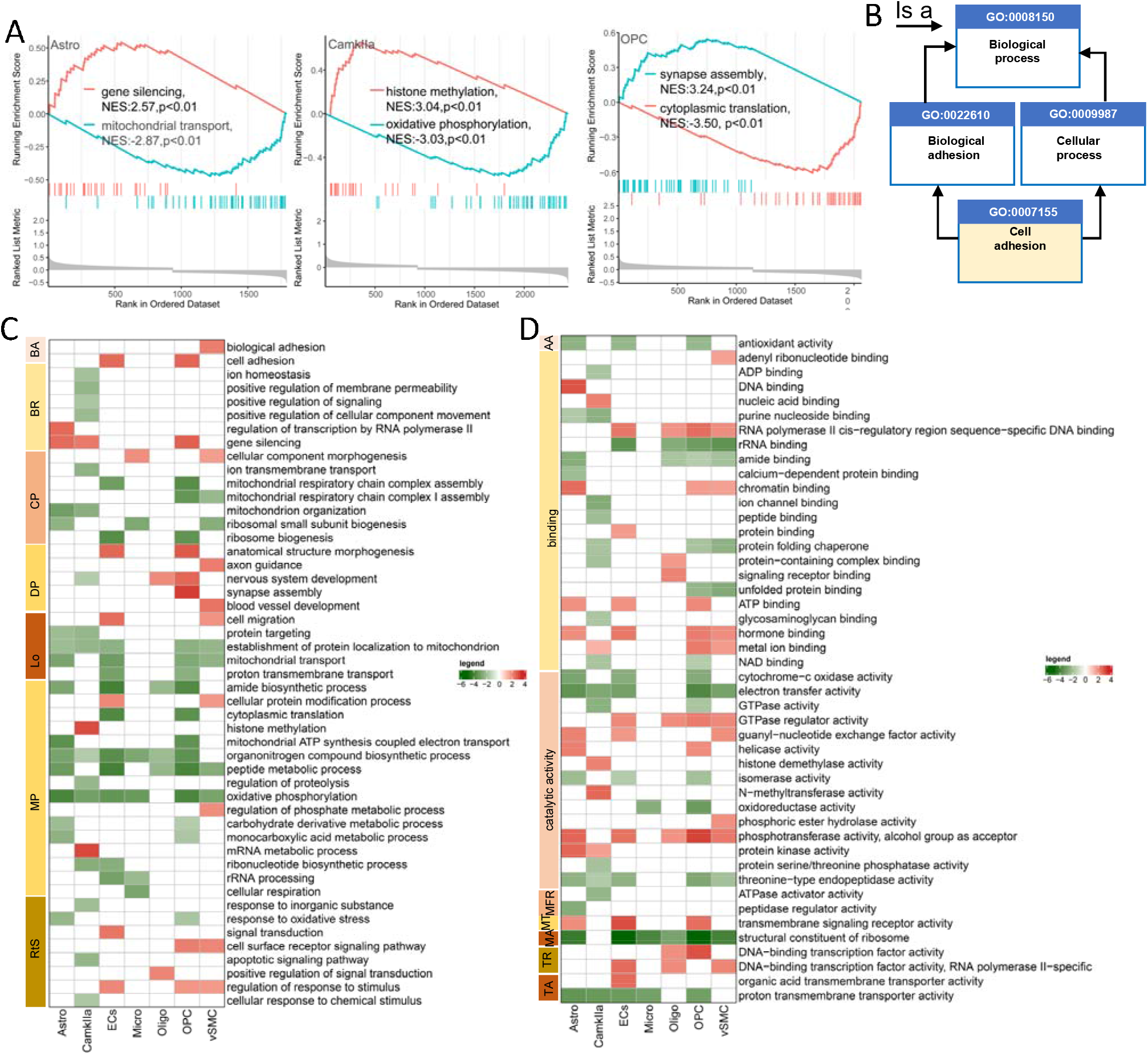
The GSEA analysis on the DEGs from different cell types. **(A)** Representative examples showing the activated and suppressed GO terms on the BP of each cell type. **(B)** The ancestor chart for the GO term (GO:0071550) cell adhesion; this term is the final child term for cellular processes and biological adhesion, their final ancestor term is biological process. **(C)** Heatmap showing the NES of the final child terms in the BP analysis. The color indicates the NES of the related term, with positive NES indicating the activated term and negative NES indicating the suppressed term. **(D)** Heatmap showing the NES of the final child terms in the MF analysis. The color indicates the NES of the related term, with positive NES indicating the activated term and negative NES indicating the suppressed term.

In addition, nerve injury induced cell type-dependent BP (**Figure 6C**) and MF changes **(Figure 6D)**. For example, in *CaMKII*α^*+*^ cells, gene silencing was probably activated via increased histone methylation and mRNA metabolic processes, indicating that epigenetic regulation occurs. Suppressive terms were detected in the BR (4 terms), Lo (2 terms), MP (4 terms) and RtS (3 terms) parts, indicating that the ion homeostasis, mitochondrial functions, proteolysis and response to stimuli are impaired due to nerve injury. In Astro and OPC, CPN ligation mainly suppressed metabolic processes (8 terms), including protein synthesis, peptide metabolic processes, oxidative phosphorylation and monocarboxylic acid metabolic process; it also inhibited the response to oxidative stress. Unlike the Astro, CPN ligation increased the DP-related process. CPN ligation also suppressed the peptide metabolic process on the Oligo (3 terms). ECs and vSMC are major cell types of brain blood vessels, and CPN ligation also suppressed nucleotide and peptide metabolic processes in ECs, which also activated the response of ECs and vSMC to stimuli and impaired protein localization in mitochondria (**Figure 6C and Table S6**).

GSEA on DEGs was mainly enriched for binding activities and catalytic activities. For *CaMKII*α^*+*^ cells, the binding to nucleic acid and peptide was suppressed, while the metal ion binding was activated. In line with the increase in histone methylation in BP analysis, histone demethylase activities were activated by CPN ligation. Activated ATP binding and hormone binding were detected on Astro, OPC, ECs and vSMC. Furthermore, the proton transmembrane transporter activity was suppressed on *CaMKII*α^*+*^, Astro, OPC Micro, and ECs (**Figure 6D and Table S6**).

### Possible changes in the ion homeostasis

Given that the same strength of peripheral stimulation activated more *CaMKII*α^+^ neurons in the RSC after CPN ligation, our scRNA-seq assay suggested that the ion homeostasis was suppressed in *CaMKIIα^+^*, which could contribute to overactivity of the RSC. To examine this possibility, we examined the expression of 36 core enriched genes from GO terms: ionic homeostasis and positive regulation of membrane permeability (**Figure S3A**). CPN ligation reduced the expression of these genes on *CaMKII*α^+^ cells and, interestingly, the *Arl6ip1*, *Atp5j*, *Atp1b1*, *Calm2*, *Fth1*, *Plp1* and *Slc25a4* were also downregulated in other cell types (**Figure S3B**). The downregulation on *Atp1a1* and *Atp1b1* indicates a possible neuronal excitability impairment in *CaMKII*^*+*^ neurons. Given that Na^+^/K^+^ -ATPase consists of two subunits, a large catalytic subunit (alpha) and a smaller glycoprotein subunit (beta), we next examined the expression of other subunits of the Na^+^/K^+^ pump: *Atp1a1-3* and Atp1b1-3. As shown in **Figure 7A**, the Na^+^/K^+^ pump subunits show a cell type-dependent expressional pattern. ECs and Micro mainly express *Atp1a1* and *Atp1b3*, while OPCs, Astro and vSMC share the same expression pattern as they mainly express *Atp1a2*, and the beta subunits include *Atp1b1* and *Atp1b2*; however, the Na^+^/K^+^ pump in the Oligo is composed of *Atp1a1* and *Atp1b3*; most *CaMKII*α^*+*^ cells expressed *Atp1a3* and *Atp1b1*, and some of them also expressed *Atp1a1* (**Figure 7B**). This cell type-dependent expression of Na^+^/K^+^ pump subunits was also presented together with the feature scatter plot in **Figure 7C**, indicating the molecular basis of resting membrane potential on different cell types. We detected downregulated *Atp1a1* in *CaMKII*α^*+*^ neurons and reduced *Atp1b1* in Astro and *CaMKII*α^*+*^ neurons (**Figure 7A**). At the same time, we observed higher *Atp1b2* in vSMC. Since Atp1a3 is also dominantly expressed in *CaMKII*α^+^ cells, changes in *Atp1a1* expression may not affect the resting membrane potential in RSC.

**Figure 7.**
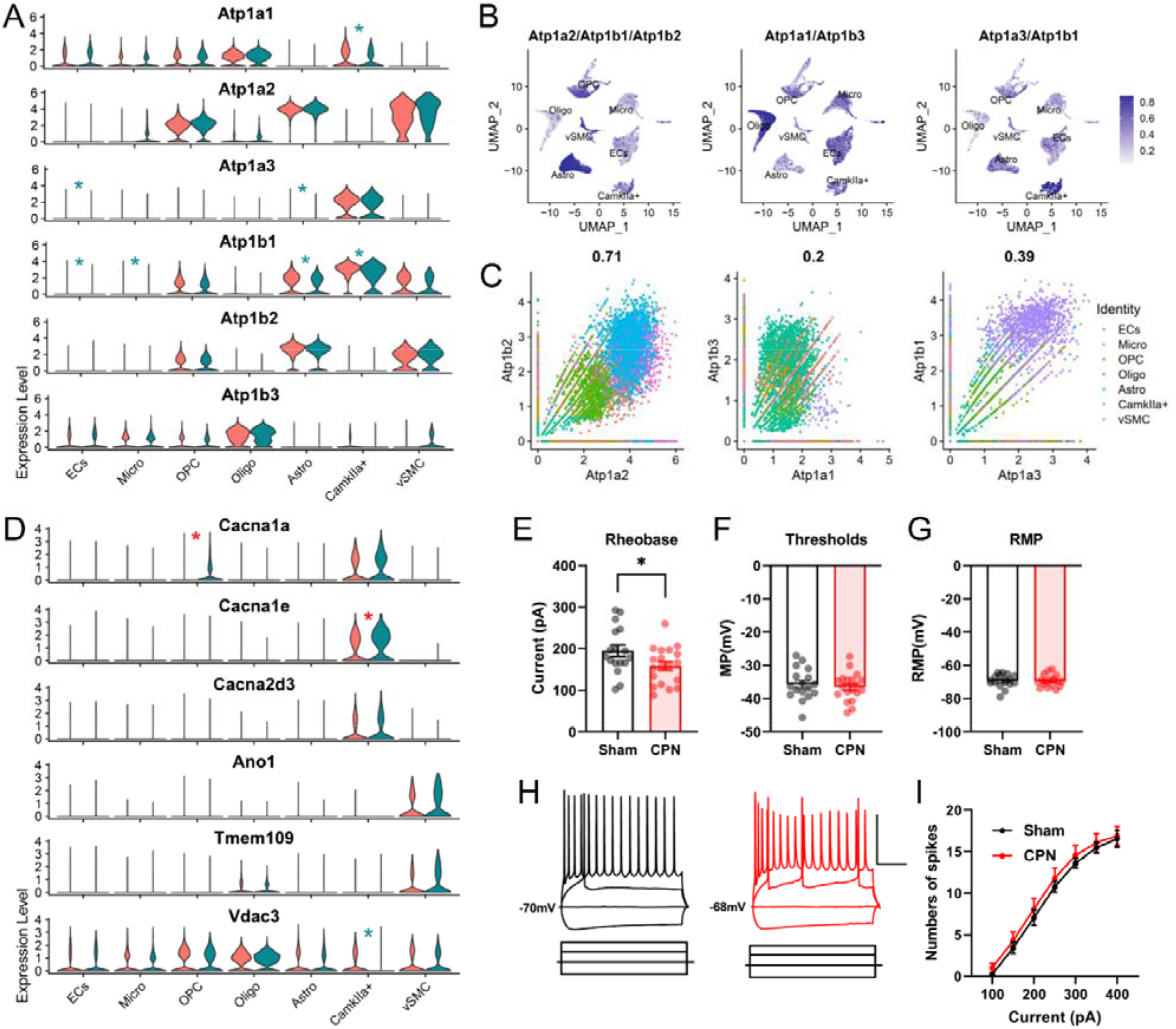
Possible effects of CPN ligation on neuronal excitability-related gene expression. (A) Violin plot showing the expression of Na^+^/K^+^ pump subunits in the RSC of mice treated with Sham (salmon) or CPN ligation (dark cyan). Cell type is dependent on the expression of Na^+^/K^+^ pump subunits. “*” in red indicates upregulation, “*” in blue indicates downregulation. (B) Umap plot showing Uscore of *Atp1a2*, *Atp1b1* and *Atp1b2* (left), *Atp1a1* and *Atp1b3* (middle), and *Atp1a3* and *Atp1b1* (right) in different cell types. (C) The feature scatter plot showing the co-expression of Na^+^/K^+^ pump subunits of *Atp1a2* and *Atp1b2* (left), *Atp1a1* and *Atp1b3* (middle), and *Atp1a3* and *Atp1b1* (right). (D) Violin plot showing cell-type dependent expression of some voltage-gated ion channels subunits in the RSC of mice treated with Sham (salmon) or CPN ligation (dark cyan). “*” in red indicates upregulation, “*” in blue indicates downregulation. (E) Whole-cell patch clamp results showing a decrease in Rheobase of neurons in the RSC by CPN ligation (Unpaired *t*-test, n = 17 for Sham, n =19 for CPN, *P* < 0.05). (F) CPN ligation does not change the firing threshold of APs recorded on RSC neurons (Unpaired *t*-test, n = 17 for Sham, n =19 for CPN, *P* > 0.05). (G) No difference in the resting membrane potentials (RMP) of pyramidal neurons in the RSC (Unpaired *t*-test, n = 17, *P* > 0.05). (H) Example showing APs that are fired under different current intensity stimulation recorded on pyramidal neurons from sham or CPN ligation group. (I) Summarized data showing that CPN ligation does not change the current-number curve of APs firing in pyramidal neurons ((Two-way ANOVA, Interaction, *F _(6, 198)_* = 0.12, *P* > 0.05; intensity, *F_(2.38, 78.81)_* = 295.1, *P* < 0.01; Sham *vs.* CPN, *F_(1, 33)_* = 0.48, *P* > 0.05, n = 17 for Sham, n = 18 for CPN).

The activities of voltage-gated ion channels (VGICs) also affect neuronal excitability. In the RSC, we examined the expression of genes encoding VGICs, most of which were enriched in *CaMKII*α^*+*^ neurons; some VGICs genes were enriched in other cell types, such as *Cacng4* in OPC and *Kcnk1* in Astro (**Figure S3C**). CPN ligation increased the expression of *Cacna1e* on *CaMKII*α^+^ cells (**Figure S3D**). We also detected upregulation of *Cacna1a* in OPC, and *Tpcn1* in Astro and vSMC. Furthermore, six genes showed CPN upregulation in vSMC, such as *Ano1* and *Tmem109*. Given that both *Cacna1e* and *Cacna2d3* encode subunits of voltage-gated calcium channels, their expression changes may impair calcium influx in *CaMKII*α^*+*^ neurons, but not neuronal excitability. To this end, we performed whole-cell patch-clamp recording on the retrosplenial neurons; we observed a slight decrease in Rheobase (**Figure 7E**), which refers to the minimal current intensity to evoke the firing of APs. Meanwhile, there were no differences in the APs firing threshold (**Figure 7F**), the resting membrane potentials (RMPs) (**Figure 7G**), as well as the input-output curve of the APs firing (**Figure 7H and I**).

### CPN ligation impairs the cell-cell communications in the RSC

Cell-cell communication is a critical part of functional cellular regulation in the brain. We next examined the possible effects of CPN ligation on cell-cell communication, including secreted signaling, extracellular matrix (ECM)-receptor interactions, and cell-cell contact interactions, via the CellChat R package^[43]^. Interestingly, a wide range of cell-cell interactions were detected in our scRNA-seq assay (**Figure 8A**). CPN ligation increased the number of interactions and the strength of the interactions (**Figure 8B**). For the incoming (**Figure 8C**) and outgoing signaling (**Figure 8D**). The cell-cell communications in the RSC showed typical cell-types dependent characteristics (**Figure 8E**). Using the selectK of CellChat R package, we identified 6 patterns of incoming patterns (**Figure 8E**) and 6 patterns of outgoing patterns (**Figure S4A**). The CPN ligation increased the strength of interactions in certain pathways, such as PDGF and NRXN pathways (**Figure 8F–J**), and decreased the strength of PTN pathway (**Figure S4B & C**); it also increased the number of interactions in the FGF, EGF and BMP pathways (**Figure 8C and D**). At the cell type level, the ECs, Oligo, Astro, Micro, and *CaMKII*α^+^ cells had fewer pathway changes than ECs and vSMC. In our scRNA-seq assay, we detected 12 upregulated pathways and 5 downregulated pathways (Table S7). The chord diagrams in **Figure 8H** and **Figure 8I** show the interaction patterns of PDGF and NRXN pathways, respectively. For the PDGF pathway, interactions from Micro to OPC, Astro and vSMC were detected in the CPN assay, but not in the Con assay (**Figure 8F**). For the NRXN pathway, CPN ligation did not change the interaction pattern (**Figure 8G**), while this pathway was increased due to higher expression of *Nrxn1-3* in OPC and Astro, which was downregulated among *CaMKII*α^+^ cells due to lower expression of *Nrxn2* and the receptor *Nlgn1-2* (**Figure 8J**).

**Figure 8.**
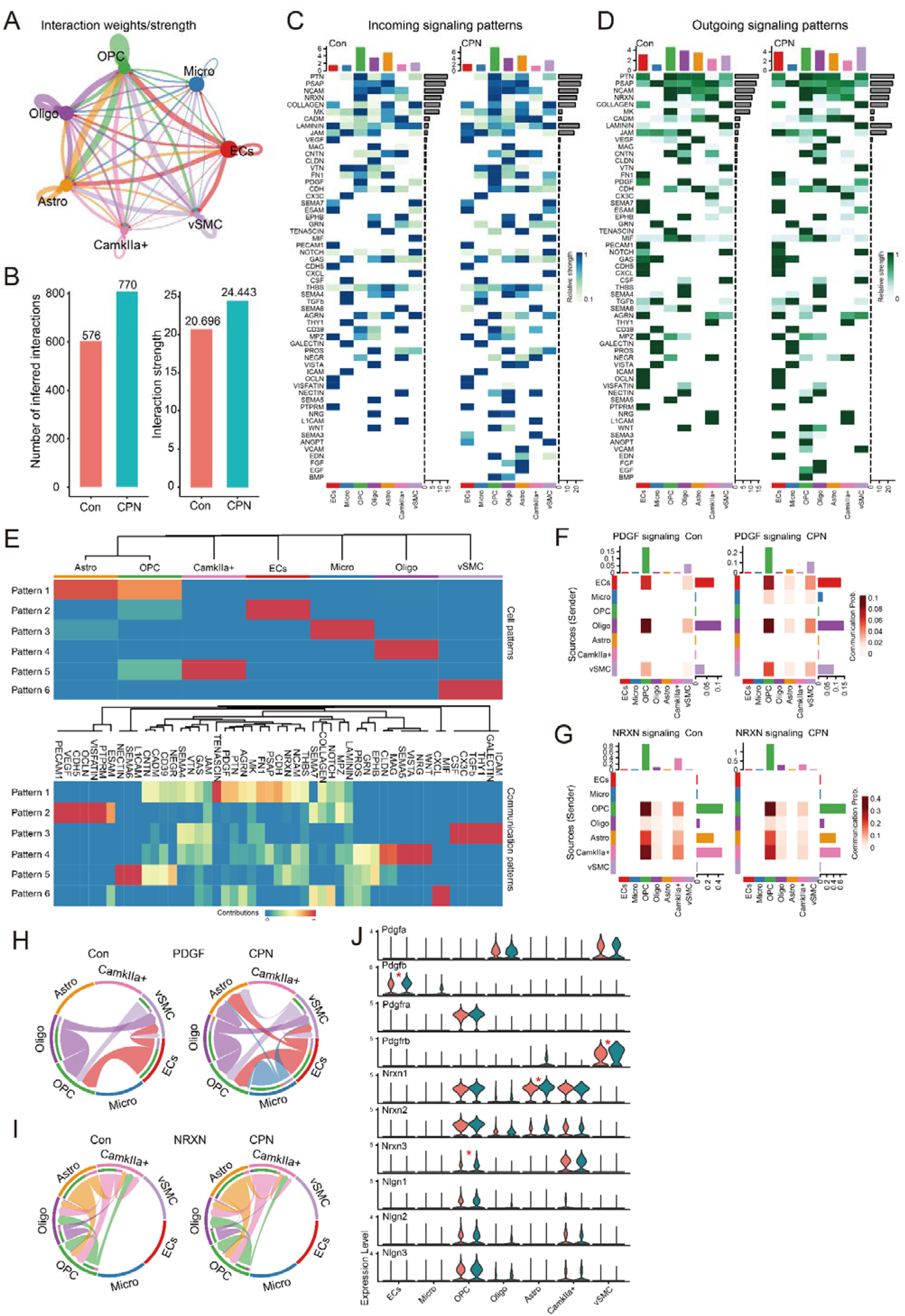
CPN ligation impairs cell-cell communication among cell types. (A) A network plot showing weighted cell-cell interactions in the RSC. (B) Histogram showing the effect of CPN ligation on the total number of cell-cell interactions and the strength of interactions in the RSC. (C) Heatmap showing the relative strengths of the identified pathways of incoming signals in the Con and CPN groups. (D) Heatmap showing the relative strengths of the identified pathways of outgoing signals in the Con and CPN groups. (E) Identify and visualize incoming communication pattern of 7 cell types from Con group. The number of patterns was determined by selectK of CellChat R package. (F) Heatmap representing the probability of PDGF pathways in the Con and CPN groups. (G) Heatmap representing the probability of NRXN pathways in the Con and CPN groups. (H) Chord diagram representing the interaction of PDGF pathways in the Con and CPN groups, with the inner bar colors indicating the receiver of signal sources from corresponding outer bars. (H) Chord diagram representing the interaction of NRXN pathways in the Con and CPN groups. (J) Violin plots showing the expression of genes in the PDGF pathway and NRXN pathway in the Con (salmon) and CPN (dark cyan) group. “*” in red indicates upregulation.

### The unbalanced excitatory/inhibitory synaptic transmission in the RSC after nerve injury

The activities of ligand-gated channels convey information transmitted from cells and regulate the excitability of cells. We examined the expression of ionotropic receptors for acetylcholine (Ach), gamma-Aminobutyric acid (GABA), glycine, glutamate, inositol 1,4,5-trisphosphate (ITP), ATP, and Ryanodine (**Figure S5A**). Interestingly, these receptors also showed cell type-dependent expression patterns, i.e., *CaMKII*α^+^ neurons expressed most of the GABA receptors, glutamate receptor subunits, as well as *Itpr1* and *Ryr3*; Oligo mainly expressed *Glrb*, *Gria2*, *Gria4*, and *Itpr2*, indicating that glutamate and glycine are its primary neurotransmitters. Mac and Micro expressed *Gria2*, *P2rx4* and *P2rx7*, and OPC and Astro also highly expressed *Gria2*, *Grid2*, *Grik5*, *Glrb* and *Itpr2*, indicating that both cell types are regulated by glutamate, glycine and ITP; moreover, both cell types expressed different subunits of GABARs and subunits of NMDARs, for example, OPC expresses *Gabrb3* and *Grin3a*, while Astro expresses *Gabra2*, *Grbrb1* and *Grin2c*; OPC also expresses *Chrna4*, which encodes the nicotinic acetylcholine receptor. The vSMC and ECs may be responsive to glutamate and Itp due to the expression of *Gria2* and ItpRs subunits, respectively. For example, vSMC expresses *Itpr1* and *Itpr2*, whereas ECs expresses *Itpr2* and *Itpr3*. In summary, the differential expression of ionotropic receptor subunits suggests that the major regulatory factors in these cells are diverse.

Next, we examined the effect of CPN ligation on the expression of ligand-gated ion channels (LGICs). In our system, CPN ligation mainly changed the cellular responses to GABA, glycine, glutamate, ITP, and Ryanodine in the RSC (**Figure 9A**). We detected higher levels of *Gria1* and *Ryr2* in *CaMKII*α^+^ neurons, as well as increased *Gria2* in OPC, Oligo, and Astro, and upregulated *Grin2c* in Astro (**Figure 9B**); these results indicate that glutamate-mediated synaptic regulations may be enhanced in *CaMKII*α^+^ neurons, OPC, Oligo, and Astro; in contrast, we detected lower levels of *Gabra1* and *Glrb* in *CaMKII*α^+^ neurons. These may indicate a decrease in inhibitory synaptic transmission in the RSC. To confirm this change in expression, we employed flow cytometer and purified AAV-CaMKIIα-mCherry-infected cells (**Figure 9 C**), and our RT-qPCR results on these purified cells also showed higher *Gria1* (**Figure 9 D**), a result that confirms the upregulation of *Gria1* in *CaMKII*α^+^ cells. we next examined the mRNA expression levels of *Gria1*, *Ryr2* from the RSC of mice treated with Sham or CPN. Similarly, we detected an increase in *Gria1* and *Ryr2*, but not *Grbra1* (**Figure 9 E**).

**Figure 9.**
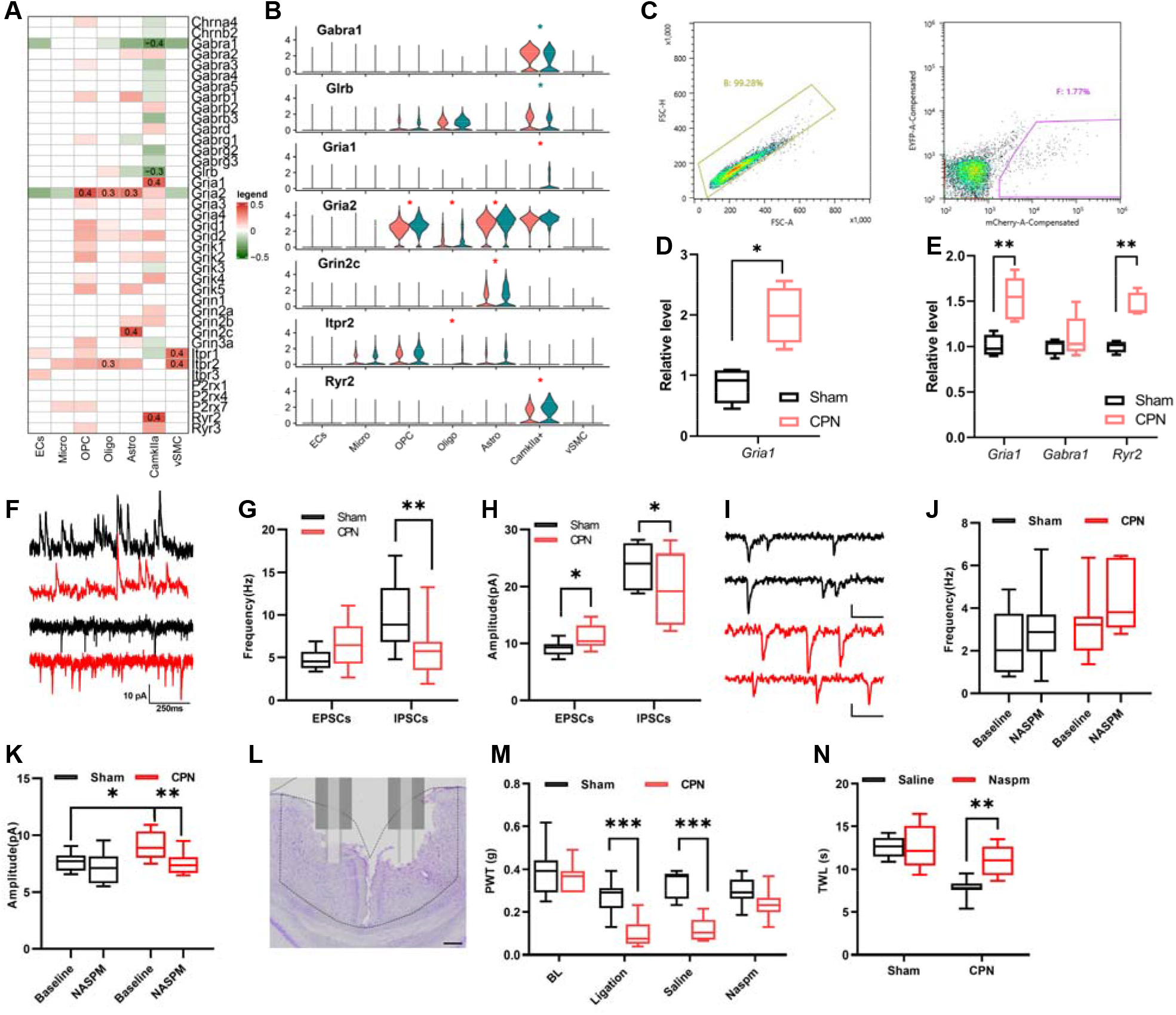
Unbalanced excitatory/inhibitory synaptic transmissions in the RSC contribute to the sensitized mechanical and thermal sensation. (A) A heatmap showing the different expressed genes coding subunits of ligand-gated receptors in the RSC after CPN ligation. The number indicates the log_2_FC of the related genes. (B) The violin plots indicate the expressions of the genes coding subunits of ligand-gated receptors the Con (salmon) and CPN (dark cyan) group. “*” in blue indicates downregulation and red reversely. (C) Flow cytometry gating strategy for *CaMKII*α^+^ neurons Analytic gating of flow cytometry data: *CaMKII*α^+^ neurons were identified from singlet cells (left); *CaMKII*α^+^ neurons were selected based on forward and side scatter (right). (D) The expressional levels of *Gria1* on the purified *CaMKII*α^+^ neurons detected via the RT-qPCR. (Unpaired *t*-test, *P* < 0.01, n = 4). (E) The expressional levels of *Gria1*, *Gabra1* and *Ryr2* in the bulk RNA from the RSC of Con and CPN groups detected via RT-qPCR. (Unpaired *t*-test, n = 4 for Sham and n = 5 for CPN, *Gria1*: *P* < 0.01; *Gabra1*: *P* > 0.05; *Ryr2*: *P* < 0.01) (F) Presented examples showing the recording of sIPSCs (up) and sEPSCs (down) under Con (Black) and CPN (Red) conditions. (G) Summarized data showing the CPN ligation reduced frequency of sIPSCs (Unpaired *t*-test, *P* < 0.05, n = 10 for Sham, n =11 for CPN), but not sEPSCs (Unpaired *t*-test, *P* > 0.05, n = 10 for Sham, n = 11 for CPN) in the RSC. (H) CPN ligation increased the amplitude of sEPSCs, but decreased the amplitude of sIPSCs in the RSC (Unpaired *t*-test, *P* < 0.05, n = 10 for Sham, n = 11 for CPN). (I) Presented examples traces of sEPSCs recorded on neurons from Sham (Black) or CPN (Red) group without (upper) or with Naspm (lower), scale bar indicates 50ms/5pA. (J) The perfusion of Naspm did not change the frequency of sEPSCs (Two-way ANOVA, Interaction, *F_(1,13)_* = 0.12, *P* = 0.74; Sham *vs.* CPN, *F_(1,13)_* = 4.87, *P* < 0.05; Baseline *vs.* Naspm, *F_(1,13)_* = 1.93, *P* > 0.05, n = 8 for Sham and n =7 for CPN). (K) The application of Naspm decreased the amplitude of sEPSCs recorded from RSC of CPN group. (Two-way ANOVA, Interaction, *F_(1,13)_* = 3.24, *P* > 0.05; Sham *vs.* CPN, *F_(1,13)_* = 4.02, *P* > 0.05; Baseline *vs.* Naspm, *F_(1,13)_* = 13.16, *P* < 0.01, n = 8 for Sham and n =7 for CPN; Sidak’s multiple comparisons test, Baseline: Sham *vs.* CPN: *P* < 0.05; Sham: Baseline vs. Naspm, *P* > 0.05; CPN: Baseline vs. Naspm, *P* < 0.01). (L) The presented example showing the locations of microinjection sites of Naspm perfusion to the RSC. (M) The microinfusing of Naspm into the RSC increased the PWTs of mice with CPN ligation. (Two-way RM ANOVA, Interaction, *F _(3, 51)_* = 6.74, *P* < 0.001; Sham *vs.* CPN, *F_(1,17)_* = 49.37, *P* < 0.001; time, *F _(2.38, 40.48)_* = 20.52, *P* < 0.01, n = 9 for Sham and n =10 for CPN, each group includes 4 female mice; Sidak’s multiple comparisons test, ligation: Sham *vs.* CPN: *P* < 0.001; Saline: Sham *vs.* CPN: *P* < 0.001; Naspm: Sham *vs.* CPN: *P* > 0.05). (N) The microinfusing of Naspm into the RSC extended the TWLs of mice with CPN ligation. (Two-way RM ANOVA, Interaction, *F _(1, 14)_* = 8.29, *P* < 0.05; Sham *vs.* CPN, *F_(1,14)_* = 24.58, *P* < 0.001; Saline vs. Naspm, *F _(1, 14)_* = 9.00, *P* < 0.01, n = 9 for Sham and n =10 for CPN, each group includes 4 female; Sidak’s multiple comparisons test, Sham: Saline *vs.* Naspm: *P* > 0.05; CPN: Saline *vs.* Naspm: *P* < 0.01).

Given that *CaMKII*α^+^ cells are distributed in different layers of the RSC, we also examined possible expression changes in different layers of the RSC. By examining the Allen Institute Brain Atlas ^[36]^, we used *Gucy1a1* and *Ddit4l* as markers for layer II/III (**Figure S5B &C**), and *Slc6a7* and *Etv1* for layer V (**Figure S5D & E**). Interestingly, an increase in *Gria1* and *Ryr2* was detected in *CaMKII*α^+^/*Slc6a7*^+^ or *CaMKII*α^+^/*Etv1*^+^ cells **(Table S8)**, indicting an increase in excitatory synaptic transmission in the deeper layers of the RSC. Furthermore, we observed a possible upregulation of *Grid2* and *Grik4* in layer II/III cells identified by *CaMKII*α^+^/ *Gucy1a1*^+^ or *CaMKII*α^+^/ *Ddit4l*^+^ cells **(Table S8)**. In summary, these observations suggest an enhanced excitatory synaptic transmission in the deeper layers of the RSC after CPN ligation.

To confirm changes in excitatory synaptic transmission, we performed whole-cell patch-clamp recordings in RSC pyramidal neurons. We recorded spontaneous excitatory postsynaptic currents (sEPSCs) and inhibitory postsynaptic currents (sIPSCs) by holding the membrane potential at −70 mV or 0 mV, respectively; besides, the application of CNQX or picrotoxin significantly blocked these currents (**Figure S5F**), indicating that these currents are mediated by AMPARs or GABARs. In line with the transcriptional changes, we observed a lower frequency of IPSCs (**Figure 9F and G**), furthermore, we also detected higher amplitude of sEPSCs but a lower amplitude of IPSCs on pyramidal neurons (**Figure 9F and H**). Therefore, CPN ligation does change the excitatory synaptic transmission in the RSC. These data indicate increased processing of input information by the RSC, which may lead to overexcitability of RSC neurons after nerve injury.

AMPARs are heterotetrametric complexes of four homologous major core subunits (GluA1-4) ^[44]^, and increasing Gria1 may increase the proportion of GluA2 lacking AMPARs, which are permeable to calcium (calcium-permeable AMPARs, CP-AMPARs), with the application of Naspm (100 μM) blocking the activities of CP-AMPARs and reducing AMPARs-mediated currents (**Figure 9I - K**). The Naspm was further microinjected into the RSC of male or female mice (**Figure 9L**) and, interestingly, this Naspm significantly elevated PWTs (**Figure 9M**) and extended the PWL of CPN ligated mice, but not of sham-operated mice (**Figure 9N**). Therefore, decreasing excitatory synaptic transmission in the RSC via Naspm changes the pain hypersensitivity induced by CPN ligation.

## Discussion

In the current study, we investigated transcriptomic changes in the RSC after peripheral nerve injury. Our data show that selective activation of *CaMKII*α^+^ neurons in the RSC decreases PWTs and induces conditioned place aversion in naïve mice. In contrast, inactivation of *CaMKII*α^+^ neurons increases PWTs in nerve-injured mice. In addition, using a single-cell RNA sequencing approach, we found that in CPN-ligated mice, cell type-dependent transcriptomic changes affect mitochondrial function, cell-cell communication, and excitatory/inhibitory synaptic transmissions, with reduced excitatory synaptic transmission via Naspm changing the CPN-ligated pain hypersensitivity in mice. Our study indicates that the RSC is a critical brain region for pain regulation, and understanding transcriptional changes in the RSC will help to understand the molecular mechanisms that maintain neuropathic pain.

### Changes in cell-cell communication in RSC

Cell-cell communication is critical for the function of specific brain regions. By taking advantage of scRNA-seq, we examined the communication patterns in RSC. Our scRNA-seq data strongly indicate that cell-cell communication patterns are cell type-dependent in the RSC (Figure 8E). CPN ligation increased the secreted signaling, such as the PDGF, VEGF, CSF and EDN pathways, the extracellular matrix, include FN1, LAMININ, TENASCIN and VTN pathways, on the Cell-Cell Contact part, the increased pathways include CDH, MPZ, NRXN and VCAM pathways. Based on connections of retrosplenial cortex ^[45–49]^, we propose that thalamic neurons may first send noxious information to *CaMKII*α^+^ neurons (**Figure 10**), and these neurons also send related information to other cell types via glutamate-mediated synaptic transmission. After nerve injury, the enhanced noxious information may activate more *CaMKII*α^+^ neurons and further induce pathological cell-cell communication between other cell types via released cytokines and growth factors, thus leading to peripheral pain sensitization. Further studies on the contributions of changed cytokines and growth factors to peripheral pain sensitivity are needed.

**Figure 10.**
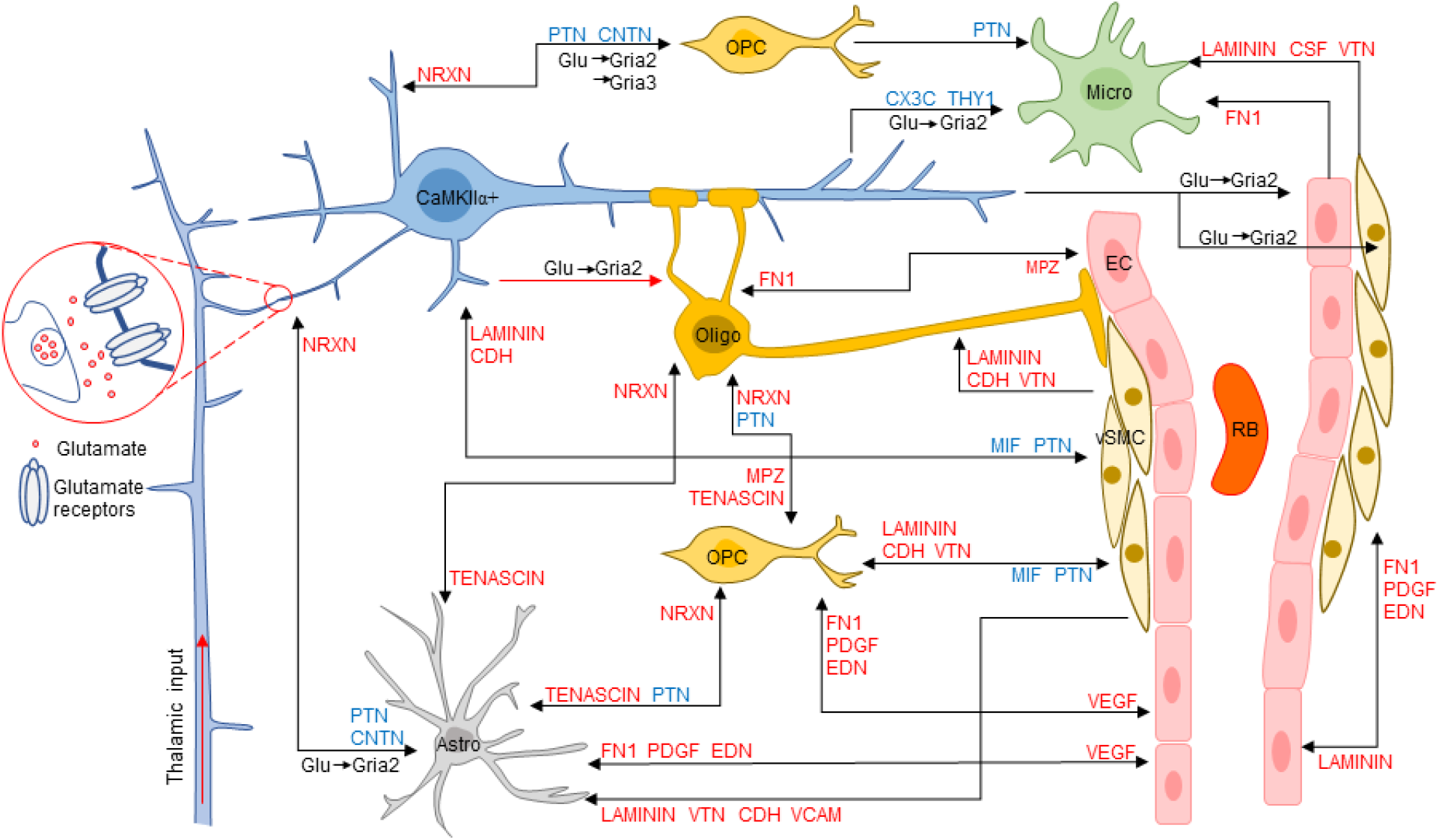
A carton demonstrating the major changes in cell-cell communication in the RSC after nerve injury. Pathway name in red indicates upregulation while in blue indicates downregulation.

The E/I synaptic balance is essential for the brain function ^[50]^. After nerve injury, the E/I imbalance usually occurs along the somatosensory pathway that leads to the persistence of chronic pain ^[51–53]^. Here, we show E/I imbalance in the RSC after nerve injury. We examined the expression of ligand-gated ion channels (LGICs) on primary neuronal transmitters, such as glutamate, glycine, GABA and ACH in the RSC. Our results show cell type-dependent expression patterns of LGICs, such as *Gria2*, which mainly expressed in *CaMKII*α^+^ neurons, OPC and Astro. In contrast, *Chnra4*, encoding the cholinergic receptor nicotinic alpha 4 subunit, was dominantly expressed in OPC, indicating its unique role in regulating OPC function. Furthermore, we observed an increase in *Gria1*, which encodes a subunit of AMPARs, the primary inotropic glutamate receptors, and a decrease in *Gabra1* and *Glrb* in *CaMKII*α^+^ neurons; these results indicate an E/I imbalance after CPN ligation. In line with this, our electrophysiological recording results confirm this and show higher amplitude of sEPSCs, but lower amplitude and frequency of sIPSCs in the RSC. This result provides information about the E/I synaptic imbalance in the RSC. It also demonstrates the usefulness of transcriptional information in understanding the molecular mechanisms of neuropathic pain.

### Neurons in the RSC regulate peripheral pain sensation

In the current study, our data reveals that enhancing the activities of *CaMKII*α^+^ neurons decreased the PWTs in naïve mice and shortened the thermal withdrawal latency. The regulation of pain sensation by the RSC lasts for less than 24 h; this implies that the regulation is not persistent. Given that noxious information was presented in the RSC, as evaluated by both electrophysiological recording ^[13]^ and brain imaging ^[54]^, these noxious responses may further facilitate peripheral pain sensation via descending regulation. Enhancing the activities of *CaMKII*α^+^ neurons in the RSC also induces place aversion. A previous study showed that formalin injection induced aversion with c-Fos expression in the RSC ^[20]^. Here we clearly show that enhancing glutamatergic neuronal activities is sufficient to induce aversion. Given that the RSC is tightly connected to the hippocampal formation ^[55,56]^ and is involved in spatial cognition ^[56–60]^, place aversion may arise from the integration of spatial information ^[15,17]^. Previous studies have shown that primary somatosensory cortex (S1), anterior cingulate cortex (ACC) ^[61]^, central amygdala ^[62]^, nucleus accumbens ^[63]^, and parabrachial nucleus ^[64]^ form a neuronal network for the regulation of pain aversion ^[65]^. Our observation suggests that the RSC is one of the brain regions of the aversion code.

Our results differ from previous studies showing that electrical stimulation of the RSC has analgesic effects on tail-flick and formalin-induced pain ^[66–69]^. Different stimulation methods can activate various neuronal assemblies in the RSC. It is well known that electrical stimulation triggers all types of cells in the RSC, including glutamatergic and GABAergic neurons. While the chemogenetic approach in the current study selectively activated *CaMKII*α^+^ neurons in the RSC, this may result in different effects on the regulation of RSC to pain sensation. In summary, we provide a piece of evidence that the activities of *CaMKII*α^+^ neurons in RSC facilitate peripheral pain sensation.

### Understanding the cellular mechanisms of neuropathic pain based on scRNA-seq results

Previous studies of DRGs in nerve-injured mice have identified the role of ATF3 in the reprograming of DRG cells. Here, we provide the first transcriptomic studies on the effects of nerve injury on RSC cells. We identified featured DEGs for each cell type, and we also examined the cell-cell communication patterns, including ECM, cell-cell contact analyzed via CellChat R package ^[43]^, growth factors, cytokines via iTalk R package ^[39]^, and major neuronal transmitters. We observed cell type-specific communication in the RSC, and these data will help to understand the coordination between retrosplenial cells.

Our data indicate that CPN ligation induces a wide range of transcriptional changes in the RSC and that most of the DEGs are cell type-dependent. For example, over 70% of DEGs are unique to the related cell types. Moreover, GO analysis on these DEGs also revealed cell type-dependent changes in BP and MF GSEA results. Considering the distinct roles of different cell types in the CNS, cell type-dependent changes may contribute to pathological changes in the brain, leading to peripheral pain hypersensitivity and comorbid anxiety or depressive symptoms. Furthermore, we observed extensive metabolic changes occurring in the RSC, and considering the energy consumption in the brain, manipulating energy support in the brain may be a novel approach to regulate pain hypersensitivity. In line with this, we also observed transcriptional changes in ECs and vSMC, which also communicate with other cell types. Unlike previous studies ^[70]^, here we emphasize cell type-dependent molecular changes in the RSC after nerve injury. In addition, the role of genetic changes in specific cell types should be further studied.

The results of scRNA-seq provide multiple clues for the molecular changes in RSC induced by peripheral nerve injury. Due to the page limitation, only changes in excitatory synaptic transmission and its role in peripheral pain hypersensitivity were confirmed here; we did not perform further confirmation of other changes. Our scRNA-seq results only present transcriptional changes in the RSC, which suggest that changes may occur at the protein level, and detailed information needs further confirmation.

## Acknowledgments

We thank Dr. San-Hua Fang at Core Facilities of the School of Medicine, Zhejiang University. The authors apologize to colleagues whose work could not be cited due to space and reference restrictions.

## Author contributions

JH Wang performed western blots, immunostaining, mechanical allodynia testing, and data analysis. YN Lian performed patch-clamp recording experiments. ZY Wang, L Liu AND L Sun carried out the immunostaining experiments and data analysis. JJ Dong, Q Wu, WJ Chen performed the virus injection and data analysis. W Chen, Z Zhang and M Zhuo designed experiments performed data analysis and approved the draft. XY Li designed experiments, performed data analysis, and wrote the paper.

## References

[1] J. Dahlhamer, J. Lucas, C. Zelaya, R. Nahin, S. Mackey, L. DeBar, R. Kerns, M. Von Korff, L. Porter, C. Helmick, MMWR. Morb. Mortal. Wkly. Rep. 2018, 67, 1001.

[2] L. Colloca, T. Ludman, D. Bouhassira, R. Baron, A. H. Dickenson, D. Yarnitsky, R. Freeman, A. Truini, N. Attal, N. B. Finnerup, C. Eccleston, E. Kalso, D. L. Bennett, R. H. Dworkin, S. N. Raja, Nat. Rev. Dis. Prim. 2017, 3, 17002.

[3] M. Costigan, J. Scholz, C. J. Woolf, Annu. Rev. Neurosci. 2009, 32, 1.

[4] M. Calvo, A. J. Davies, H. L. Hébert, G. A. Weir, E. J. Chesler, N. B. Finnerup, R. C. Levitt, B. H. Smith, G. G. Neely, M. Costigan, D. L. Bennett, Neuron 2019, 104, 637.

[5] S. Tang, H. Jing, Z. Huang, T. Huang, S. Lin, M. Liao, J. Zhou, J. Cell. Biochem. 2020, 121, 1635.

[6] R. Kuner, T. Kuner, Physiol. Rev. 2021, 101, 213.

[7] R. R. Ji, C. R. Donnelly, M. Nedergaard, Nat. Rev. Neurosci. 2019, 20, 667.

[8] M. Tsuda, Neurosci. Bull. 2018, 34, 178.

[9] C.-L. Li, K.-C. Li, D. Wu, Y. Chen, H. Luo, J.-R. Zhao, S.-S. Wang, M.-M. Sun, Y.-J. Lu, Y.-Q. Zhong, X.-Y. Hu, R. Hou, B.-B. Zhou, L. Bao, H.-S. Xiao, X. Zhang, Cell Res. 2016, 26, 83.

[10] G. Hu, K. Huang, Y. Hu, G. Du, Z. Xue, X. Zhu, G. Fan, Sci. Rep. 2016, 6, 1.

[11] J. K. Niehaus, B. Taylor-Blake, L. Loo, J. M. Simon, M. J. Zylka, Neuron 2021, 109, 1274.

[12] J.-A. Yang, J.-M. He, J.-M. Lu, L.-J. Jie, J. Cell. Biochem. 2018, 119, 4792.

[13] R. W. Sikes, L. J. Vogt, B. A. Vogt, Pain 2008, 135, 160.

[14] G. C. Quintero, J. Pain Res. 2013, 6, 713.

[15] G. Vantomme, Z. Rovó, R. Cardis, E. Béard, G. Katsioudi, A. Guadagno, V. Perrenoud, L. M. J. Fernandez, A. Lüthi, Cell Rep. 2020, 31, DOI 10.1016/j.celrep.2020.107747.

[16] J. P. Shine, J. P. Valdés-Herrera, M. Hegarty, T. Wolbers, J. Neurosci. 2016, 36, 6371.

[17] A. S. Alexander, D. A. Nitz, Curr. Biol. 2017, 27, 1551.

[18] J. T. Da Silva, D. A. Seminowicz, Pain reports 2019, 4, e732.

[19] P. E. Paulson, T. J. Morrow, K. L. Casey, Pain 2000, 84, 233.

[20] L. G. Lei, Y. Q. Zhang, Z. Q. Zhao, Neuroreport 2004, 15, 67.

[21] C. E. Kim, Y. K. Kim, G. Chung, H. J. Im, D. S. Lee, J. Kim, S. J. Kim, Neuroimage 2014, 86, 311.

[22] B. A. Vogt, The Cingulate Cortex in Neurologic Diseases: History, Structure, Overview, Elsevier B.V., 2019.

[23] T. V. P. Bliss, G. L. Collingridge, B. K. Kaang, M. Zhuo, Nat. Rev. Neurosci. 2016, 17, 485.

[24] C. S. Chiou, C. C. Chen, T. C. Tsai, C. C. Huang, D. Chou, K. Sen Hsu, Anesthesiology 2016, 125, 779.

[25] K. I. Vadakkan, Y. H. Jia, M. Zhuo, J. Pain 2005, 6, 747.

[26] S. R. Chaplan, F. W. Bach, J. W. Pogrel, J. M. Chung, T. L. Yaksh, J. Neurosci. Methods 1994, 53, 55.

[27] Y. J. Wang, Z. X. Zuo, C. Wu, L. Liu, Z. H. Feng, X. Y. Li, Front. Mol. Neurosci. 2017, DOI 10.3389/fnmol.2017.00289.

[28] M. Leger, A. Quiedeville, V. Bouet, B. Haelewyn, M. Boulouard, P. Schumann-Bard, T. Freret, Nat. Protoc. 2013, 8, 2531.

[29] K. Hargreaves, R. Dubner, F. Brown, C. Flores, J. Joris, Pain 1988, 32, 77.

[30] K. J. Livak, T. D. Schmittgen, Methods 2001, 25, 402.

[31] G. J. Brewer, J. R. Torricelli, Nat. Protoc. 2007, 2, 1490.

[32] S. Zhong, W. Ding, L. Sun, Y. Lu, H. Dong, X. Fan, Z. Liu, R. Chen, S. Zhang, Q. Ma, F. Tang, Q. Wu, X. Wang, Nature 2020, 577, 531.

[33] D. Aran, A. P. Looney, L. Liu, E. Wu, V. Fong, A. Hsu, S. Chak, R. P. Naikawadi, P. J. Wolters, A. R. Abate, A. J. Butte, M. Bhattacharya, Nat. Immunol. 2019, 20, 163.

[34] T. Stuart, A. Butler, P. Hoffman, C. Hafemeister, E. Papalexi, W. M. Mauck, 3rd Y. Hao, M. Stoeckius, P. Smibert, R. Satija, Cell 2019, 177, 1888.

[35] N. Gehlenborg, 2019.

[36] E. S. Lein, M. J. Hawrylycz, N. Ao, M. Ayres, A. Bensinger, A. Bernard, A. F. Boe, M. S. Boguski, K. S. Brockway, E. J. Byrnes, L. Chen, L. Chen, T.-M. Chen, M. C. Chin, J. Chong, B. E. Crook, A. Czaplinska, C. N. Dang, S. Datta, N. R. Dee, A. L. Desaki, T. Desta, E. Diep, T. A. Dolbeare, M. J. Donelan, H.-W. Dong, J. G. Dougherty, B. J. Duncan, A. J. Ebbert, G. Eichele, L. K. Estin, C. Faber, B. A. Facer, R. Fields, S. R. Fischer, T. P. Fliss, C. Frensley, S. N. Gates, K. J. Glattfelder, K. R. Halverson, M. R. Hart, J. G. Hohmann, M. P. Howell, D. P. Jeung, R. A. Johnson, P. T. Karr, R. Kawal, J. M. Kidney, R. H. Knapik, C. L. Kuan, J. H. Lake, A. R. Laramee, K. D. Larsen, C. Lau, T. A. Lemon, A. J. Liang, Y. Liu, L. T. Luong, J. Michaels, J. J. Morgan, R. J. Morgan, M. T. Mortrud, N. F. Mosqueda, L. L. Ng, R. Ng, G. J. Orta, C. C. Overly, T. H. Pak, S. E. Parry, S. D. Pathak, O. C. Pearson, R. B. Puchalski, Z. L. Riley, H. R. Rockett, S. A. Rowland, J. J. Royall, M. J. Ruiz, N. R. Sarno, K. Schaffnit, N. V Shapovalova, T. Sivisay, C. R. Slaughterbeck, S. C. Smith, K. A. Smith, B. I. Smith, A. J. Sodt, N. N. Stewart, K.-R. Stumpf, S. M. Sunkin, M. Sutram, A. Tam, C. D. Teemer, C. Thaller, C. L. Thompson, L. R. Varnam, A. Visel, R. M. Whitlock, P. E. Wohnoutka, C. K. Wolkey, V. Y. Wong, M. Wood, M. B. Yaylaoglu, R. C. Young, B. L. Youngstrom, X. F. Yuan, B. Zhang, T. A. Zwingman, A. R. Jones, Nature 2007, 445, 168.

[37] G. Yu, L. G. Wang, Y. Han, Q. Y. He, Omi. A J. Integr. Biol. 2012, 16, 284.

[38] S. Jin, C. Guerrero-Juarez, L. Zhang, I. Chang, P. Myung, M. Plikus, Q. Nie, 2020, 1.

[39] Y. Wang, R. Wang, S. Zhang, S. Song, C. Jiang, G. Han, M. Wang, J. Ajani, A. Futreal, L. Wang, bioRxiv 2019, 507871.

[40] William Revelle, 2021.

[41] C. Katche, J. H. Medina, Cereb. Cortex 2017, 27, 1060.

[42] D. A. Barrière, A. M. Hamieh, R. Magalhães, A. Traoré, J. Barbier, J. M. Bonny, D. Ardid, J. Busserolles, S. Mériaux, F. Marchand, Pain 2019, 160, 2241.

[43] S. Jin, C. F. Guerrero-Juarez, L. Zhang, I. Chang, R. Ramos, C.-H. Kuan, P. Myung, M. V Plikus, Q. Nie, Nat. Commun. 2021, 12, 1088.

[44] S. F. Traynelis, L. P. Wollmuth, C. J. McBain, F. S. Menniti, K. M. Vance, K. K. Ogden, K. B. Hansen, H. Yuan, S. J. Myers, R. Dingledine, Pharmacol. Rev. 2010, 62, 405.

[45] T. van Groen, B. A. Vogt, J. M. Wyss, Neurobiol. Cingulate Cortex Limbic Thalamus 1993, 123.

[46] T. Van Groen, J. M. Wyss, J. Comp. Neurol. 2003, 463, 249.

[47] N. Yamawaki, K. A. Corcoran, A. L. Guedea, G. M. G. Shepherd, J. Radulovic, 2019, 2728.

[48] T. van Groen, J. M. Wyss, J. Comp. Neurol. 1990, 300, 593.

[49] T. van Groen, J. M. Wyss, J. Comp. Neurol. 1992, 315, 200.

[50] N. Z. Gungor, J. Johansen, Neuron 2019, 102, 903.

[51] F. Le Cao, M. Xu, K. Gong, Y. Wang, R. Wang, X. Chen, J. Chen, J. Pain 2019, 20, 917.

[52] D. Vecchia, D. Pietrobon, Trends Neurosci. 2012, 35, 507.

[53] M. Petrou, R. Pop-Busui, B. R. Foerster, R. A. Edden, B. C. Callaghan, S. E. Harte, R. E. Harris, D. J. Clauw, E. L. Feldman, Acad. Radiol. 2012, 19, 607.

[54] G. Wik, H. Fischer, B. Bragée, M. Kristianson, M. Fredrikson, Neuroreport 2003, 14, 619.

[55] J. Sugar, M. P. Witter, N. M. van Strien, N. L. M. Cappaert, Front. Neuroinform. 2011, 5, DOI 10.3389/fninf.2011.00007.

[56] N. Yamawaki, K. A. Corcoran, A. L. Guedea, G. M. G. Shepherd, J. Radulovic, Cereb. Cortex 2019, 29, 2728.

[57] A. N. Opalka, D. V. Wang, Learn. Mem. 2020, 27, 310.

[58] S. D. Vann, J. P. Aggleton, E. A. Maguire, Nat. Rev. Neurosci. 2009, 10, 792.

[59] A. M. P. Miller, L. C. Vedder, L. M. Law, D. M. Smith, Front. Hum. Neurosci. 2014, 8, 586.

[60] A. S. Mitchell, R. Czajkowski, N. Zhang, K. Jeffery, A. J. D. Nelson, Brain Neurosci. Adv. 2018, 2, 239821281875709.

[61] X. B. Wu, L. N. He, B. C. Jiang, X. Wang, Y. Lu, Y. J. Gao, Neurosci. Bull. 2019, 35, 613.

[62] G. Corder, B. Ahanonu, B. F. Grewe, D. Wang, M. J. Schnitzer, G. Scherrer, Science (80-.). 2019, 363, 276.

[63] R. Al-Hasani, J. G. McCall, G. Shin, A. M. Gomez, G. P. Schmitz, J. M. Bernardi, C. O. Pyo, S. Il Park, C. M. Marcinkiewcz, N. A. Crowley, M. J. Krashes, B. B. Lowell, T. L. Kash, J. A. Rogers, M. R. Bruchas, Neuron 2015, 87, 1063.

[64] M. C. Chiang, A. Bowen, L. A. Schier, D. Tupone, O. Uddin, M. M. Heinricher, J. Neurosci. 2019, 39, 8225.

[65] R. Du, W. J. Luo, K. W. Geng, C. L. Li, Y. Yu, N. Wei, J. Chen, Neurosci. Bull. 2020, 36, 649.

[66] G. M. Reis, R. S. Fais, W. A. Prado, Pharmacol. Biochem. Behav. 2015, 131, 112.

[67] G. M. Reis, A. C. Rossaneis, J. W. S. Silveira, Q. M. Dias, W. A. Prado, J. Pain 2011, 12, 523.

[68] G. M. Reis, Q. M. Dias, J. W. S. Silveira, F. Del Vecchio, N. Garcia-Cairasco, W. A. Prado, J. Pain 2010, 11, 1015.

[69] A. C. Rossaneis, G. M. Reis, W. A. Prado, Pharmacol. Biochem. Behav. 2011, 100, 220.

[70] W. Renthal, I. Tochitsky, L. Yang, Y. C. Cheng, E. Li, R. Kawaguchi, D. H. Geschwind, C. J. Woolf, Neuron 2020, 108, 128.

