## Supplemental figure 1-5 for "Single-cell RNA sequencing uncovers the excitatory/inhibitory synaptic unbalance in the retrosplenial cortex after peripheral nerve injury"

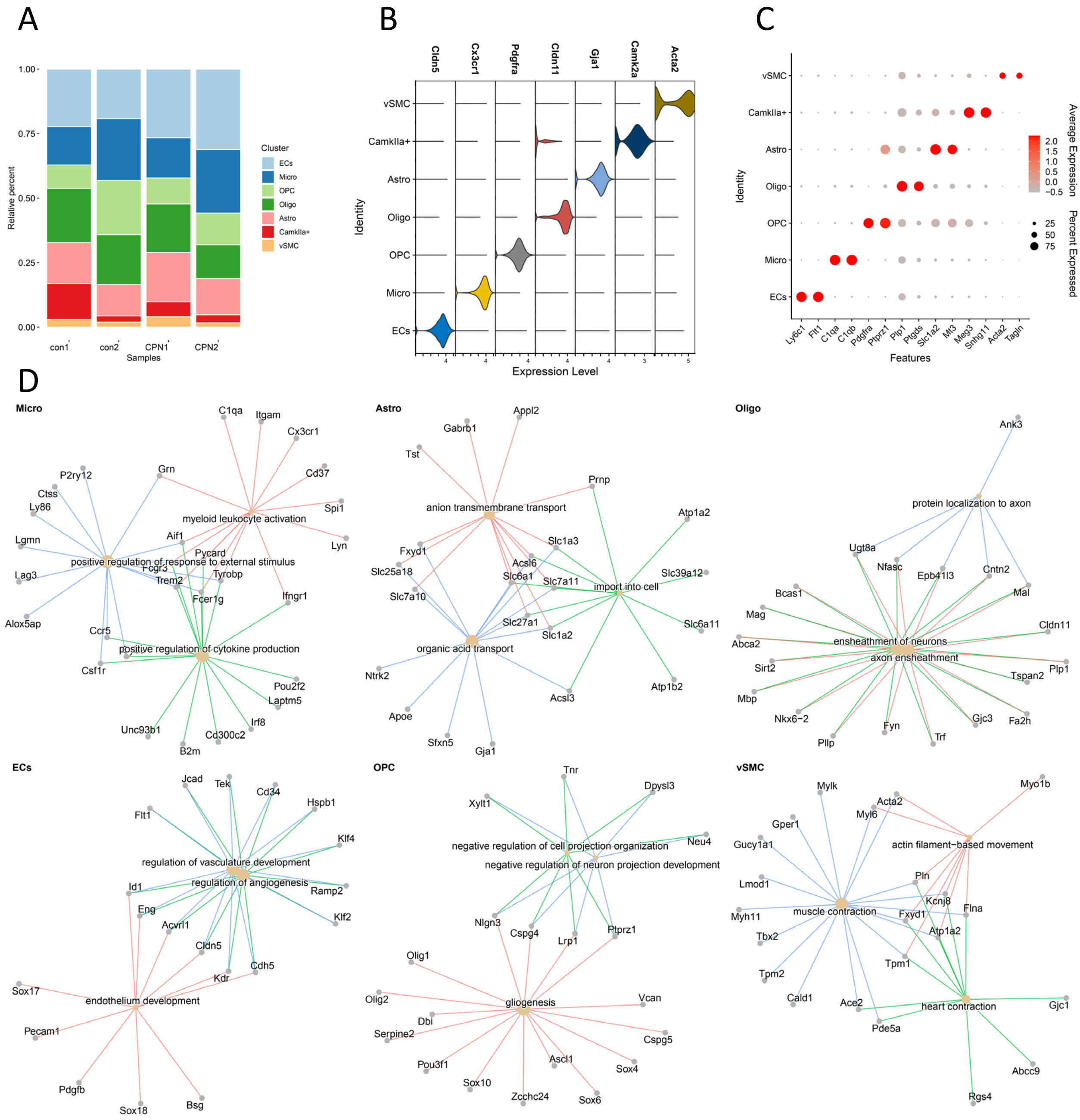


**Figure S1｜The cell types of RSC with or without CPN ligation.**

(A) Bar chart of the relative percent of 7 cell types in 4 samples.

(B) Violin plot showing expression level of classical cell type markers in different cell types.

(C) Dot plot showed top 2 fDEGs for each cell type.

(D) fDEGs related to typical biological process in each cell type besides CaMKIIα$^{+}$ neurons.

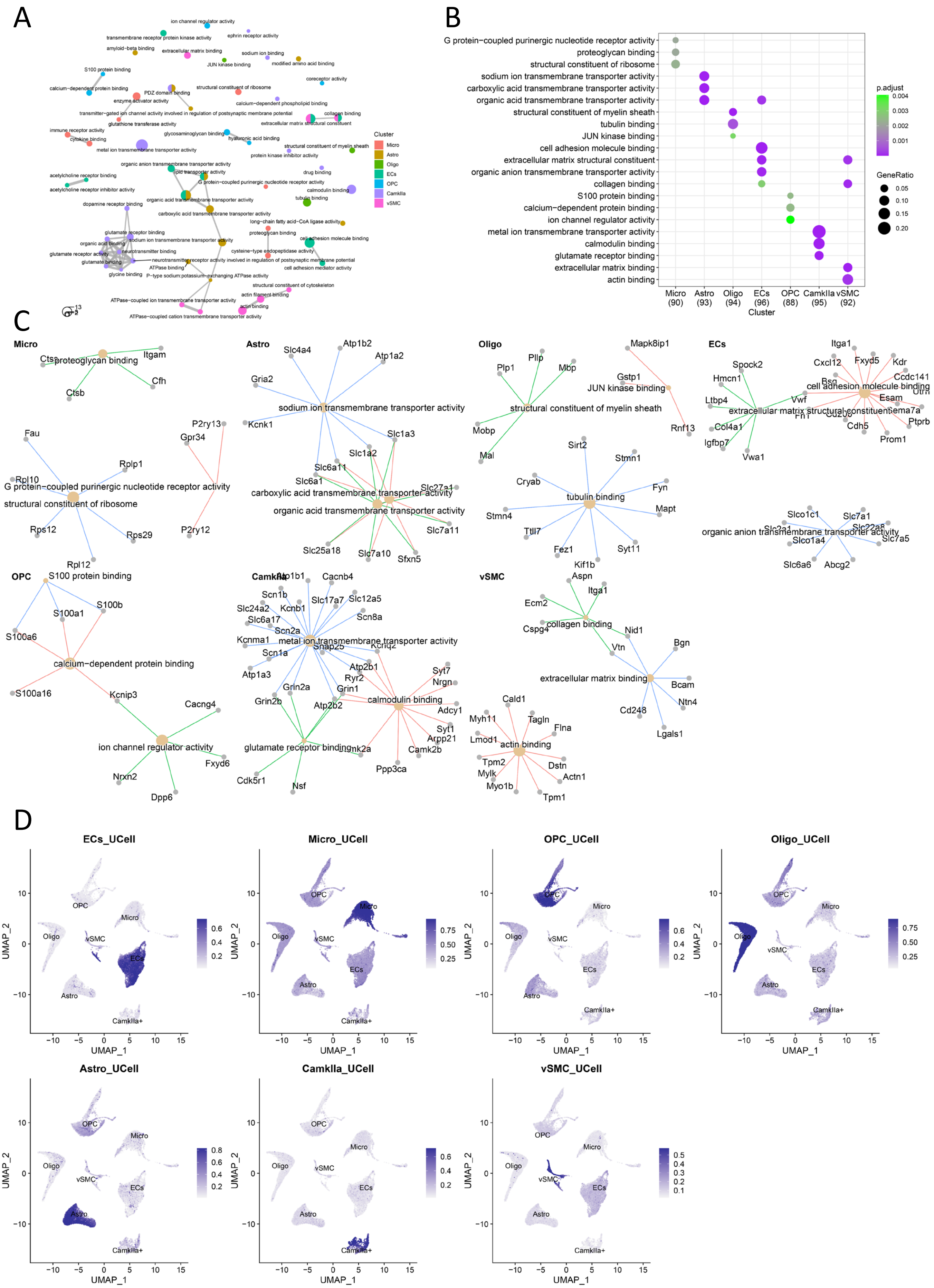
A
B
C
D
Cluster
Micro
Astro
Oligo
ECs
OPC
CamkIIa
vSMC
p.adjust
GeneRatio
Micro (90)
Astro (93)
Oligo (94)
ECs (96)
OPC (88)
CamkIIa (95)
vSMC (92)
G protein−coupled purinergic nucleotide receptor activity
proteoglycan binding
structural constituent of ribosome
sodium ion transmembrane transporter activity
carboxylic acid transmembrane transporter activity
organic acid transmembrane transporter activity
structural constituent of myelin sheath
tubulin binding
JUN kinase binding
cell adhesion molecule binding
extracellular matrix structural constituent
organic anion transmembrane transporter activity
collagen binding
S100 protein binding
calcium−dependent protein binding
ion channel regulator activity
metal ion transmembrane transporter activity
calmodulin binding
glutamate receptor binding
extracellular matrix binding
actin binding
ECs_UCell
Micro_UCell
OPC_UCell
Oligo_UCell
Astro_UCell
CamkIIa_UCell
vSMC_UCell
UMAP_1
UMAP_2

**Figure S2 | Networks showing the Go analysis of molecular functions in different cell types of RSC.**

(A) Molecular function network derived from top 100 fDEGs of different cell types.

(B) Dot plot showing the GO analysis on the molecular function of top 100 fDEGs

(C) fDEGs related to typical molecular function in each cell type.

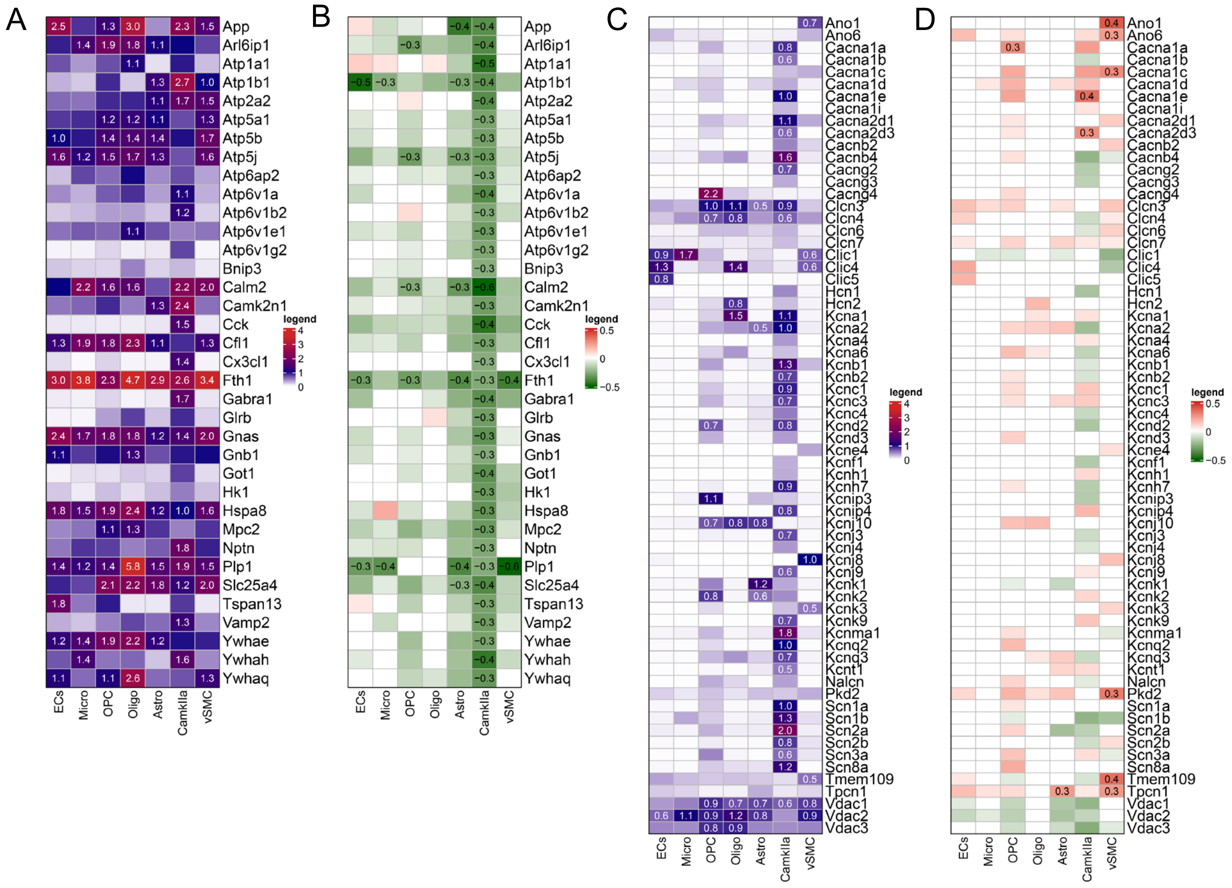

**Figure S3 | Heatmaps showing the expressional changes of the ion channels related genes.**

(A) Heatmap showing the cell-type dependent expression level of 36 core enriched genes from GO terms: ionic homeostasis and positive regulation of membrane permeability

(B) Heatmap showing the cell-type dependent expression fold-change of 36 core enriched genes from GO terms: ionic homeostasis and positive regulation of membrane permeability

(C) Heatmap showing the cell-type dependent gene expression level of voltage-gated ion channels.

(D) Heatmap showing the cell-type dependent gene expression fold-change of voltage-gated ion channels in CPN group comparing to Con group.

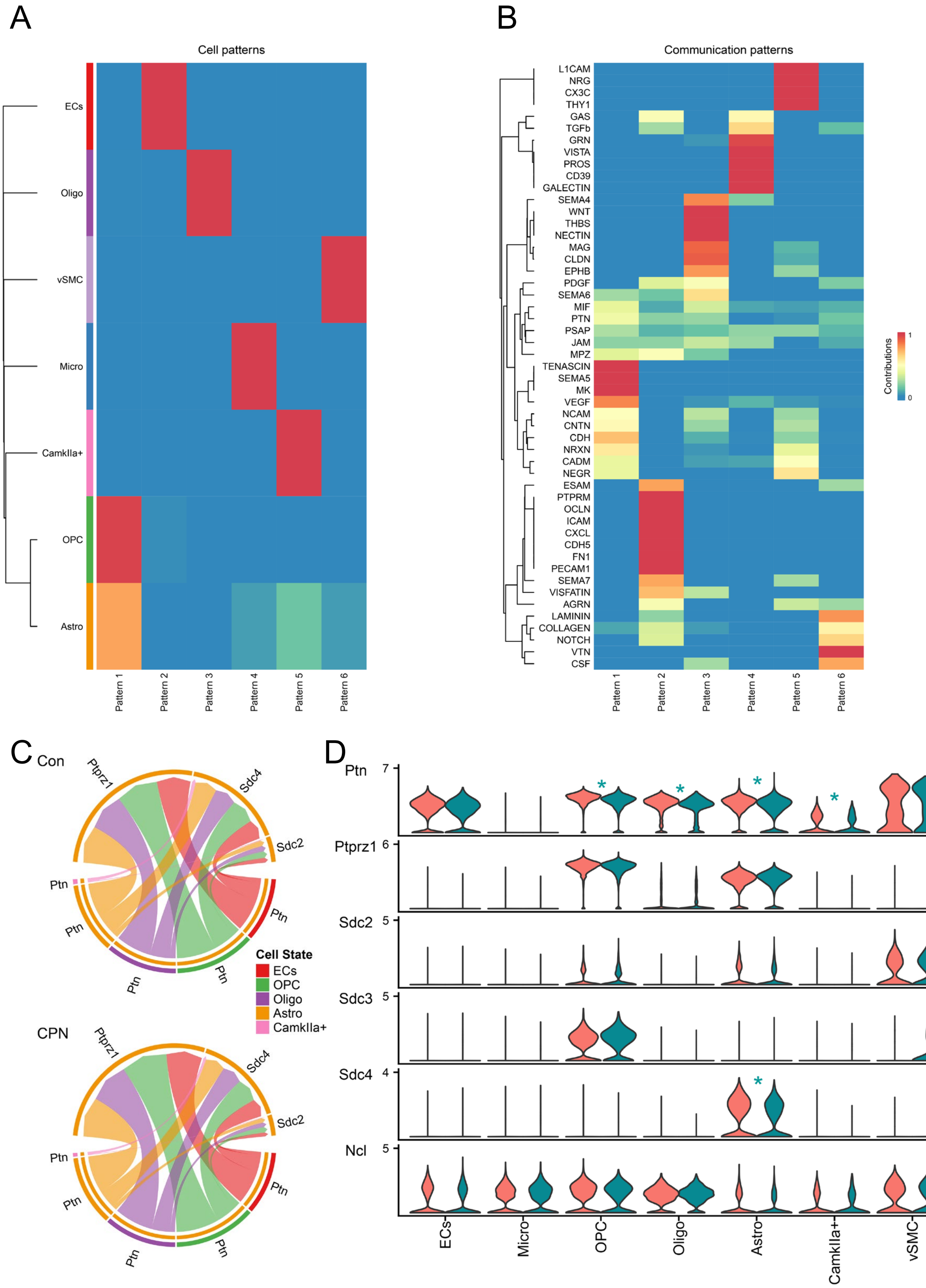


**Figure S4 | The possible changes on the cell-cell communications.**

(A) Identify outgoing communication pattern of 7 cell types from Con group. The number of patterns was determined by selectK of CellChat package.

(B) visualize outgoing communication pattern of 7 cell types from Con group.

(C) Chord plot showed the PTN signaling pathway network in Con and CPN group.

(D) Violin plot showed the gene expression distribution of PTN pathway. * in blue indicates downregulation.

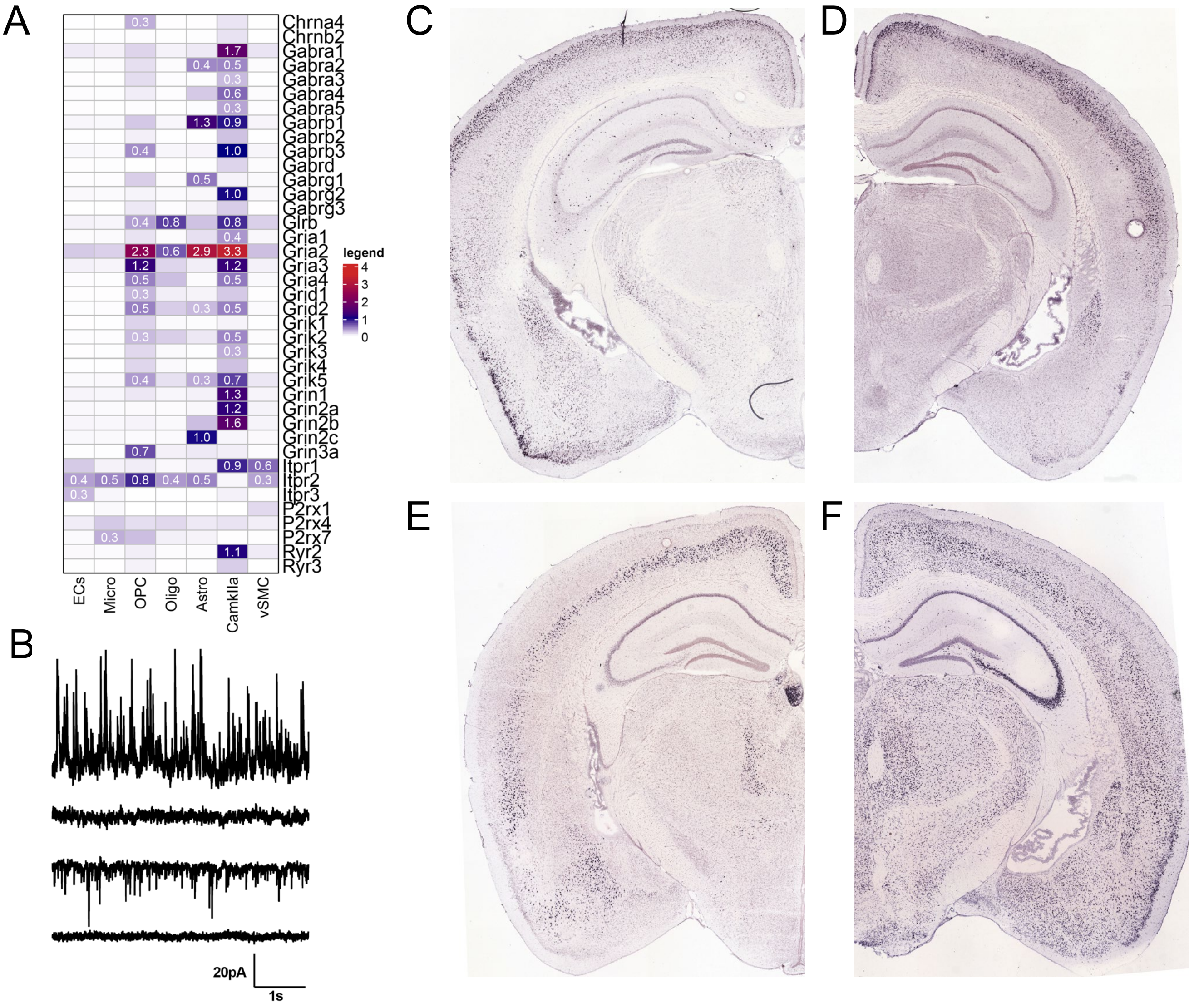


**Figure S5 | The gene expression pattern of ligand-gated channels .**

(A) Heatmap showing the cell-type dependent gene expression level of ligand-gated channels.

(B) Presented examples showed that the spontaneous currents recorded at 0 mV (upper 2 traces) and -70 mV (lower 2 traces) were respectively blocked by picrotoxin and CNQX.

(C) *Gucy1a3* (*Gucy1a1*) mainly expressed in layer II/III of cerebral cortex (https://mouse.brain-map.org/experiment/show/77866848)

(D) *Ddit4l* mainly expressed in layer II/III of cerebral cortex. (https://mouse.brain-map.org/experiment/show/71836878)

(E) *Etv1* mainly expressed in layer V of cerebral cortex. (https://mouse.brain-map.org/experiment/show/72119595)

(F) *Slc6a7* mainly expressed in layer V of cerebral cortex. (https://mouse.brain-map.org/gene/show/88708). Fig C-F was modified from Allen Mouse Brain Atlas, available from https://mouse.brain-map.org (Lein et al., 2007).

**Title of the supplementary tables.**

**Table S1. The featured differential expression genes (fDEGs) of each cell types.**

**Table S2. The Go enrichments analysis on the BP of the top 100 fDEGs in different cell types.**

**Table S3. The Go enrichments analysis on the MF of the top 100 fDEGs in different cell types.**

**Table S4. The DEGs between the Con and CPN group in different cell types and Further analysis of common or unique upregulated and downregulated DEGs.**

**Table S5. The Gene set enrichment analysis on the BP of DEGs in different cell types.**

**Table S6. The Gene set enrichment analysis on the MF of DEGs in different cell types.**

**Table S7. Summary of upregulated and downregulated pathways in Cell-cell communication analysis.**

**Table S8. Genes expression in layer II/III and layer V CaMKIIα$^{+}$ neurons.**
